# S100A1ct: a synthetic peptide derived from human S100A1 protein improves cardiac contractile performance and survival in pre-clinical heart failure models

**DOI:** 10.1101/2023.03.04.531024

**Authors:** Dorothea Kehr, Julia Ritterhoff, Manuel Glaser, Lukas Jarosch, Rafael E. Salazar, Kristin Spaich, Karl Varadi, Jennifer Birkenstock, Michael Egger, Erhe Gao, Walter J. Koch, Hugo A. Katus, Norbert Frey, Andreas Jungmann, Cornelius Busch, Paul J. Mather, Arjang Ruhparwar, Mirko Völkers, Rebecca C. Wade, Patrick Most

## Abstract

**Background:** The EF-hand Ca^2+^ sensor protein S100A1 has been identified as a molecular regulator and enhancer of cardiac performance. S100A1’s ability to recognize and modulate the activity of targets such as SERCA2a and RyR2 in cardiomyocytes has mostly been ascribed to its hydrophobic C-terminal *α*-helix (residues 75-94). **Objective:** We therefore hypothesized that a synthetic peptide consisting of residues 75-94 of S100A1 and an N-terminal solubilization tag (S100A1ct) could mimic the performance enhancing effects of S100A1 and may be suitable as a peptide therapeutic to improve the function of diseased hearts. **Methods and Results:** Applying an integrative translational research pipeline, ranging from computational molecular modeling to large animal cardiac disease models, we characterize S100A1ct as a cell-penetrating peptide with positive inotropic and antiarrhythmic properties in normal and failing myocardium *in vitro* and *in vivo*. This activity translates into improved contractile performance and survival in pre-clinical heart failure models with reduced ejection fraction after S100A1ct systemic administration. Mechanistically, S100A1ct exerts a fast and sustained dose-dependent enhancement of cardiomyocyte Ca^2+^ cycling and prevents ß-adrenergic receptor triggered Ca^2+^ imbalances by targeting SERCA2a and RyR2 activity. Modeling suggests that S100A1ct may stimulate SERCA2a by interacting with the sarcoplasmic transmembrane segments of the multi-span integral membrane Ca^2+^ pump. Incorporation of a cardiomyocyte targeting peptide tag into S100A1ct (cor-S100A1ct) further enhanced its biological and therapeutic potency *in vitro* and *in vivo*. **Conclusion:** S100A1ct peptide is a promising lead for the development of a novel peptide-based therapeutic against heart failure with reduced ejection fraction.

## Introduction

Peptide therapeutics have played a notable role in medical practice since the advent of insulin therapy in the 1920s. Currently, there are more than one hundred US Food and Drug Administration (FDA) and European Medical Agency (EMA) approved peptide medicines on the market.^1^ Peptidic drugs are being increasingly appreciated due to their on-target selectivity, specificity, potency and safety. A reason is that polypeptides and their targets have co-evolved to such extent that it is rather difficult to design an artificial molecular entity, such as a small molecule, that substantially mimics the accuracy of fit and coverage of peptidic structures at their target sites.^2, 3^ Not surprisingly, a robust pipeline with 155 peptide drugs in clinical trials and more than 500 peptide-based candidates in preclinical development against a wide range of intracellular and extracellular targets may result in a broad clinical use across all medical disciplines in the years to come, including cardiovascular diseases.^2, 3^

It is against this background that we report the development of the S100A1ct peptide as a potential therapeutic agent for systemic treatment against heart failure with reduced ejection fraction (HFrEF). S100A1ct is a short synthetic peptide comprising the C-terminal helix (amino acids; aa 75-94) of the native human S100A1 protein and a 6-residue N-terminal tag to aid chemical synthesis and solubility (Figure 1A). The parent protein S100A1 (1-94 aa) is a homodimer and a member of the EF-hand calcium (Ca^2+^) sensor protein superfamily, which comprises at least 20 paralogs.^4, 5^ S100A1 is expressed in a tissue- and cell-specific manner with highest abundance in cardiac and skeletal muscle, where it resides at the sarcoplasmic reticulum (SR), the sarcomere and within the mitochondria of myocytes.^6^ Numerous *in vitro* and *in vivo* genetic gain- and loss-of function studies by us and others have shown that S100A1 plays a decisive role as a molecular enhancer of heart and skeletal muscle contractile performance.^7^ This is due in part to the ability of S100A1 to bind to and regulate the activity of a number of key molecular effectors including the SR Ca^2+^ ATPase (SERCA2a), the ryanodine receptor (RyR), mitochondrial ATP synthase (F1-ATPase) and titin, all of which govern cardiac and skeletal muscle Ca^2+^ cycling, energy supply and mechanical properties.^4, 6, 7^

**Figure 1.**
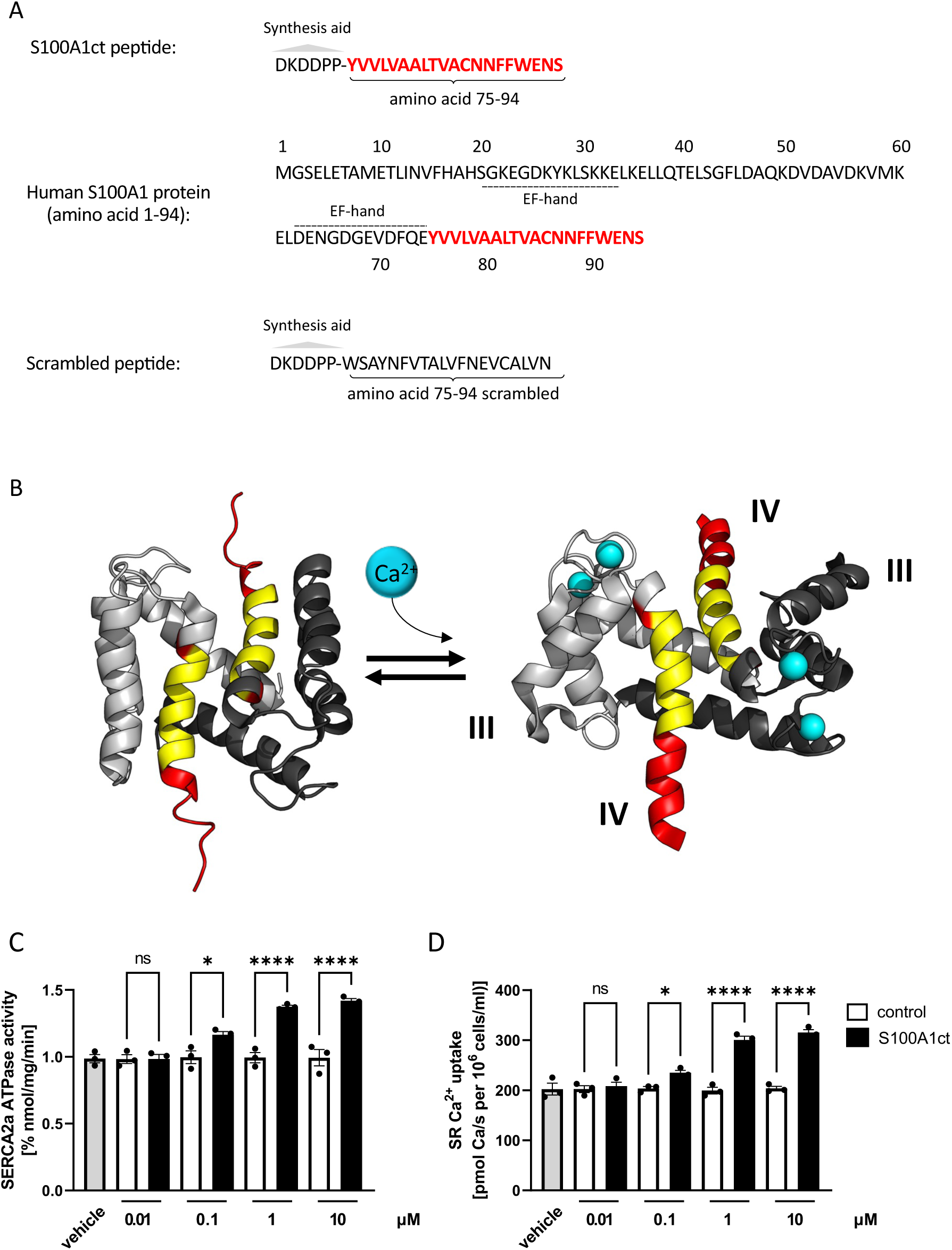
A, Primary sequences of the S100A1ct peptide (top) and the parent human S100A1 protein (NCBI reference NP_006262) (middle). The C-terminal helix (amino acids; aa 75-94) is highlighted in red both in the S100A1ct and S100A1 protein sequence. The dotted lines indicate S100A1’s two EF-hand Ca^2+^ binding sites. The scrambled control peptide sequence is shown at the bottom. An N-terminal short peptide tag (DKDDPP; grey triangle) aided the scalable production and solubility of the peptides. B, 3D structures of Ca^2+^ free (apo, PDB-ID: 2L0P, NMR model 1) and Ca^2+^ bound (holo, PDB-ID: 2LP3, NMR model 1; Ca^2+^ ions are shown as cyan spheres) human S100A1 protein (shown in gray cartoon representation with different shades of gray indicating the two subunits of the S100A1 dimer). The exposed C-terminal domain of holo-S100A1 (aa 75-94, i.e., helix IV) is highlighted in red, except for the apolar sequence stretch ‘VVLVAALTVA’, which is shown in yellow (the respective region is also shown in the same color-coding for apo-S100A1).C, Dose-dependent augmentation of SERCA2a ATPase activity by 0.01, 0.1, 1 and 10 μM S100A1ct peptide in cardiac SR vesicle preparations compared to control. Measurements were carried out in triplicates. Data are given as mean±SEM for SERCA2a activity at a free Ca^2+^ assay concentration of 1 μM. *P=0.0208; ****P<0.001. D, Dose-dependent enhancement of SERCA2a-mediated SR Ca^2+^ uptake by 0.01, 0.1, 1 and 10 μM S100A1ct peptide in saponin-permeabilized rabbit ventricular cardiomyocytes compared to control treatment. Measurements were performed in duplicate. Data are given as mean±SEM for SR Ca^2+^ uptake at a free Ca^2+^ assay concentration of 1 μM. *P=0.0158; ****P<0.001. Control: scrambled peptide, vehicle: aqueous carrier used for scrambled and S100A1ct peptide dilution.

Nuclear magnetic resonance (NMR) and crystallographic determination of S100A1’s three-dimensional (3D) structure and dynamics indicated that a hydrophobic binding pocket becomes exposed upon Ca^2+^ binding (Figure 1B), providing an explanation for the Ca^2+^-dependent interactions of S100A1 with its intracellular targets.^8–10^ Computational modeling identified the hydrophobic residues in the alpha-helical C-terminus of S100A1 as able to interact, with e.g., its respective molecular epitopes on the RyR.^11^ Consistently, comparative molecular assays utilizing chemically permeabilized skeletal muscle fibers demonstrated an equipotent enhancement of RyR1-dependent SR Ca^2+^-release and isometric force development by human recombinant S100A1 protein and by the S100A1ct peptide.^12^ Further experiments, using chemically permeabilized ventricular cardiomyocytes, showed a similar rate of attenuation of diastolic RyR2 Ca^2+^ spark frequency by the S100A1 protein and the S100A1ct peptide, as well as the peptide’s ability to mimic not only the activated “on-state” of the parent protein but to outcompete S100A1’s binding at the RyR2 receptor.^13^

These *in vitro* results provide the starting point for the systematic *in vitro* and *in vivo* investigation of the biological and therapeutic properties of S100A1ct that we report here, resulting in the clinically relevant discovery that the S100A1ct peptide, owing to its cell-penetrating ability, is a promising lead for the development of peptide-based therapeutics against HFrEF. This was achieved by using a vertical translational research approach spanning from systematic investigations and modelling of S100A1ct actions in molecular model systems to *in vitro* and *in vivo* S100A1ct treatment studies in failing human and rodent cardiomyocytes from HFrEF hearts and in mouse and pig HFrEF models, respectively.

Thus, our translational study, starting from the S100A1ct peptide’s structure, dynamics and molecular interactions, and extending to *in vivo* activity in small and large animals will provide a basis for future studies that may enable S100A1ct peptide based safe and efficacious HFrEF treatment in a first-in-human study with a convenient dosing regimen.

## Materials and Methods

Additional resources are available from the corresponding author upon reasonable request. A more detailed description of each method and protocol is provided in the supplement. All animal procedures and experiments were carried out according to the ‘Guide for the Care and Use of Laboratory Animals’ (National Institutes of Health) and were approved by the local Institutional Animal Care and Use Committee of Baden-Württemberg, Germany and Jefferson University, PH, USA. All mice were housed at 22°C with a 12-hour light, 12-hour dark cycle with free access to water and standard chow. Bay cage types were used for housing pigs. Per pig, a body weight prescribed floor space of at least 0.5 to 0.7 m^2^ was provided according to directive 2010/63/EU with straw bedding. The environment was enriched by pellet balls, chains and gnawing rods. Humidity and temperature were kept at 50-60 % and 20-24 °C, respectively. Access to water was unlimited and restricted food provided twice/day (SAF130M).

### Generation of human recombinant S100A1 protein and S100 peptides

Human recombinant S100A1 protein (hrS100A1) was generated and purified as described previously.^14, 15^ Synthetic S100 and control peptides were custom-made by commercial suppliers (Eurogentec, GeneScript and Bachem) and sequences are given in Figure 1A, S1B and 5A.

### Calcium ATPase activity and sarcoplasmic reticulum calcium uptake assays

The activity of SERCA2a and SR Ca^2+^ uptake was determined as previously reported by the use of SR vesicle preparations^12^ and saponin-skinned rabbit left ventricular cardiomyocytes, respectively.^15, 16^ SR vesicles were prepared from murine hearts of homozygous S100A1-knock out mice as given elsewhere.^17^

### F1-ATPase activity

The ATPase activity of purified bovine F1-ATPase was measured at 37°C with an ATP-regenerating system as reported by following the oxidation of NADH to NAD+ at 340 nm in a Hewlett-Packard spectrophotometer as described previously.^18, 19^

### Isolation and culture of ventricular cardiomyocytes

Enzymatic isolation of adult rat left ventricular (LV) cardiomyocytes (AVCMs) was performed as described elsewhere.^20^ Normal rat AVCMs were isolated from hearts of euthanized healthy male Sprague-Dawley (SD) rats while failing rat AVCMs were obtained from male SD rats with post-myocardial infarction heart failure with reduced ejection fraction as previously detailed.^21^ Human failing cardiomyocyte isolation and culture was performed as given elsewhere.^22^

### Myofilament calcium-force relationship assay

Measurements of the calcium-force relationship in Triton-skinned rabbit ventricular trabeculae (average length/width/thickness: 3,500 ± 300/153 ± 30/124 ± 27 μm; sarcomere length adjusted to 2.2 μm set by laser diffraction) were done as described previously.^15, 23^

### Biomimetic parallel artificial membrane permeability assay

The membrane permeability of S100A1ct peptide and hrS100A1 protein was assessed by using a commercially purchased parallel artificial membrane permeability assay kit and suppliers protocol (BioAssays Systems; Catalog No : PAMPA-096).

### Confocal microscopic imaging

Confocal microscopic transmission and fluorescent images of FITC-labeled S100A1ct and control treated cardiomyocytes were taken as described previously.^13^

### Computational analysis of S100 peptide sequences

Peptide secondary structure was analyzed with PEP2D.^24^ The FMAP 2.0 server^25^ was used to predict the positioning and energetics of transmembrane alpha-helices in input peptide sequences.

### Computational modeling of S100A1ct/SERCA2a complexes

A structure of S100A1ct was extracted from molecular dynamics simulations^26^ in which the N-terminal tag was unstructured and the rest of the peptide was α-helical. S100Act was then docked to structures of human SERCA2a E1 (PDB-ID: 6JJU)^27^ and E2 (PDB-ID: 7BT2)^28^ states using a customized two-step procedure that was validated against experimentally determined structures for other peptides that bind SERCA. Briefly (see further details in supplemental methods), the ClusPro 2.0 webserver^29^ was used with default settings and the “Balanced” scoring function for global rigid-body docking of the peptide and SERCA2a. Then, the docked poses were post-processed to account for the presence of the membrane using RosettaMP tools by refining and re-ranking the docked structures using MPDock.^29, 30^

### Cellular calcium transient analysis, sarcoplasmic reticulum Ca^2+^ load and Ca^2+^ leak assessment

FURA2-AM based epifluorescent measurement of calcium transients in the various AVCM preparations were performed as described elsewhere.^21, 22, 31^

### Cellular Ca^2+^ spark analysis

Ca^2+^ sparks in intact quiescent AVCMs were monitored using a Leica SP2, (Mannheim, Germany) laser scanning confocal microscope (LSCM) as described previously.^13^ Recorded Ca^2+^ sparks were quantified using an automated algorithm adapted from a previously published method. ^32^

### Cellular contraction analysis

Contractile properties of isolated AVMC preparations were obtained by video-edge-detection, as described previously.^21, 33^ Analysis of steady-state twitches at indicated stimulation rates, 37°C, and 2 mM [Ca^2+^]_extracellular_ was performed by custom-designed software written in LabView (version 5.0; National Instruments).

### Immunoblotting

The phosphorylation status of serine-22/23 troponin I and serine-16 phospholamban in homogenates of AVCM preparations was assessed as described previously with purchased phospho-site specific (Cell signaling: Phospho-Troponin I (Cardiac) (Ser23/24) antibody #4004; Badrilla: PLB pSer16 antibody Catalogue No.: A010-12) and total protein (Cell signaling: Troponin I Antibody #4002 and Phospholamban (D9W8M) Rabbit mAb #14562) antibodies.

### Murine post-myocardial infarction heart failure model

HFrEF was induced by surgical ligation of the left anterior descending artery in male C57/B6 mice as previously described.^34^

### Porcine post-myocardial infarction cardiac dysfunction model

Cardiac systolic contractile dysfunction in German farm pigs was induced by percutaneous transluminal catheter-based balloon occlusion of the left circumflex coronary artery as previously described.^31^

### Echocardiographic and hemodynamic analysis of cardiac Function

Transthoracic cardiac echocardiography in mice was performed as described previously.^35^ Cardiac contractile performance in mice and farm pigs was obtained by LV pressure-tip and pressure-volume catheterization as reported elsewhere.^35, 36^

### Clinical chemistry, blood count and electrocardiogram

Clinical chemistry organ biomarker and basic blood count from murine and porcine blood, plasma and serum samples was conducted by Thomas Jefferson Hospital laboratory services and Heidelberg University Hospital clinical chemistry unit. Porcine electrocardiograms were measured and recorded as described elsewhere.^37^

### Statistical analysis

The numbers of independent experiments/ animals are specified in the relevant figure legends. Data are expressed as mean ± standard error of the mean (SEM). Statistical analysis was performed with Prism 8.0 or 9.0 software (GraphPad). Normal distribution of data was verified by Shapiro-Wilk test. For normal distributed data, statistical comparisons between 2 groups were conducted by unpaired, two-tailed t-test. Statistical comparisons between 3 or more groups were conducted by one-way or two-way ANOVA followed by a Tukey posthoc analysis or Šídák’s multiple comparisons test to determine statistical significance. For non-normal distributed data, a Mann-Whitney test was performed for comparisons between 2 groups and a Kruskal-Wallis test was performed between 3 or more groups with subsequent Dunn’s test for multiple testing. For comparison over time, Bonferroni posthoc analysis was performed. The value of p < 0.05 was considered statistically significant.

## Results

### Molecular effects of S100A1ct peptide on effector proteins that regulate cardiomyocyte performance regulation

As previous molecular studies demonstrated the equivalent effect of the S100A1ct peptide and the Ca^2+^-activated S100A1 protein on diastolic RyR2 function,^13^ we hypothesized that S100A1ct may impact further target proteins of S100A1 in cardiomyocytes, such as SERCA2a or mitochondrial F1-ATP synthase. Figure 1A depicts the primary sequence of the synthetic S100A1ct peptide derived from the human S100A1 gene. Due to its high hydrophobicity, a hydrophilic N-terminal 6-mer tag was included in the sequence to assist the chemical synthesis and solubility in aqueous carrier solutions. A synthetic peptide with the same aa composition in a random order was attached to the 6-mer tag and used as a scrambled control (Figure 1A). Figure 1B illustrates the position of the respective C-terminal *α*-helix IV within the quaternary structure of holo-S100A1 protein. To systematically assess the effect of S100A1ct on SERCA2a and F-ATPase synthase activity, we performed the same molecular assays that were previously used ^12, 17–19^ to investigate the functional impact of recombinant human S100A1 (rhS100A1) protein on these proteins. Figure 1C depicts the S100A1ct dose-dependent increase of SERCA2a activity by S100A1ct using sarcoplasmic reticulum (SR) vesicles that were prepared from homozygous S100A1 knock-out murine hearts. Moreover, incubation of purified F1-ATPase protein with increasing amounts of S100A1ct yielded a dose-dependent enhancement in F1-ATP synthase activity (Figure S1A), as previously demonstrated for rhS100A1 protein.^18, 19^ To corroborate the specificity of these findings, the corresponding 20 aa synthetic C-terminal fragments of the S100 protein paralogs S100A4 (S100A4ct, NP_002952.1) and S100B (S100Bct, NP_006263.1) with the same N-terminal 6-residue tag were compared to S100A1ct (Figure S1B). Neither S100A4ct nor S100Bct impacted the activity of SERCA2a and F1-ATPase (Figure S1C and S1D). The results of our molecular assays were further strengthened by the observation of enhanced SR Ca^2+^ sequestration in chemically skinned permeabilized rabbit ventricular cardiomyocytes by S100A1ct (Figure 1D), as previously been shown for rhS100A1 protein.^15, 16^ Consistent with other published data reporting a decrease in the Ca^2+^ sensitivity of myofilament of chemically skinned rabbit ventricular trabeculae due to hrS100A1 protein,^15, 23^ S100A1ct diminished the Ca^2+^ sensitivity in the same multicellular assay system to the same extent (data not shown).

### Modeling of the S100A1ct/SERCA2a transmembrane interaction

To gain a better understanding of how S100A1ct might affect SERCA2a activity, we next generated *in-silico* models of the putative S100A1ct/SERCA2a complex. Secondary structure predictions with PEP2D indicated that S100A1ct has a pronounced helical region around its apolar sequence stretch ‘VVLVAALTV’ and some helicity in its C-terminal part (see Supplementary Figure S2A). Our previous molecular dynamics simulations of S100A1ct in aqueous environment^26^ corroborate the high helicity for the apolar sequence stretch and show that the C-terminal part is transiently helical and able to undergo conformational switching. Nevertheless, from the simulations, the energetically most stable conformation of the S100A1ct-20mer is, apart from the unstructured tag, fully α-helical in a polar environment. Furthermore, S100A1ct is predicted by FMAP 2.0 to form a transmembrane helix (see Supplementary Table S1 and Supplementary Figure S3A), similarly to phospholamban (PLN) and sarcolipin (SLN) (see Supplementary Table S1 and Supplementary Figure S3D and E, respectively), two known predominantly α-helical (Supplementary Fig S2D and E, respectively) and transmembrane oligopeptide inhibitors of SERCA,^38^ although its membrane interaction is predicted to be much less energetically favorable than for PLN/SLN (see Supplementary Table S1). Strikingly, the S100Bct and S100A4ct peptides and the scrambled control peptide, which had no effect on SERCA2a activity, are not predicted to exist as stable transmembrane α-helices (see Supplementary Table S1 and Supplementary Figures S2 and S3). We used our customized docking pipeline for generating putative structures of the complexes of S100A1ct with SERCA2a in the E1 and E2 states (see Supplementary Table S2). A customized pipeline was necessary because standard docking procedures do not account for the membrane environment and the first step of our docking pipeline led to poses predominantly docking to the SERCA2a transmembrane domain. Similar docking patterns were obtained for the E1 and the E2 states of SERCA2a. We identified 5 possible binding sites for the peptide on SERCA2a (see Supplementary Figure S4): the groove lined by the transmembrane helices TM2/TM6/TM9, which is the site at which PLN and SLN bind in experimentally determined structures of SERCA (cf. PDB-IDs 4Y3U^39^ or 4H1W^40^); a groove lined by TM3/TM5/TM7 on the opposite side of SERCA2a from the PLN-binding site; and, less prominently, the site formed by helices TM1/TM3/TM4, the site formed by the helices TM1 and TM2 (the “cytoplasmic bordering” helix), and the site formed by helices TM8/TM9/TM10. The S100A1ct peptide mostly adopted an orientation with its helix axis perpendicular to the membrane plane or slanted similarly to the helices in SERCA2a itself (see Supplementary Figure S4). Its orientation facilitated interactions between the residues in the S100A1ct helix and the transmembrane helices of SERCA2a. Docking poses were obtained with either the N-terminus of the S100A1ct peptide pointing towards the cytoplasmic side of the membrane (up-orientation; the same orientation as observed in experimentally determined structures of PLN /SLN) or the N-terminus of the S100A1ct peptide pointing towards the lumenal side of the membrane (down orientation) (see Supplementary Figure S4). Overall, the two orientations had similar docking scores (see Supplementary Table S2), however, the up orientation was preferred for the PLN-binding site (see Figure 2A and Supplementary Figure S5D). A clear tendency for the down orientation was observed for the TM3/TM5/TM7-binding site of the E1 state at the top-ranking positions after MPDock refinement (see Figure 2A), while for the E2 state, the up orientation was at rank 1 (see Supplementary Figure S5A), followed by the down orientation at rank 2 and 3 (Supplementary Figure S5B and S5C). For the E1 state, a pose in the TM1/TM3/TM4 site was also obtained in which a disulfide bridge could be formed between C86 of the S100A1ct and C841 of SERCA2a (see Figure 2A). In this pose, the S100A1ct also tended to slightly kink at the “CNN” part of the sequence (see Figure 2A). For the E1 state, the PLN-binding site was ranked higher (rank 2) compared to the E2 state (rank 4), where the TM3/TM5/TM7-binding site was favored at the top-3 ranks (see Figure 2A vs Figure S5). Considering the possibility that the re-ranking score for the docking pose to the E1 state featuring disulfide bridge formation was dominated by this effect, the PLN-binding site can even be considered at rank 1 for docking to the E1 state. Interestingly, a patch of apolar residues lining the PLN-binding site in the E1 state looks less compact for the E2 state (cf. Figure 2A vs Supplementary Figure S5D) and the peptide was shifted a bit out of the PLN-binding site after the refinement step (Supplementary Figure S5D vs Supplementary Figure 4F – red pose), indicating that it might be “unhappy” at this site, which could not be observed for the peptide docked to the E1 state (cf. Figure 2A vs Supplementary Figure 4A – pink pose). The TM1/TM2 site (e.g., see Supplementary Figure S4B) directly borders (and also encompasses) the TM2 helix of the PLN-binding site and thus it might function as an accessory or intermediate binding site for it. Based on our docking results, a possible hypothesis for the SERCA2a activation effect of S100A1ct might be by perturbing binding of the inhibitory peptide PLN to SERCA2a.

**Figure 2.**
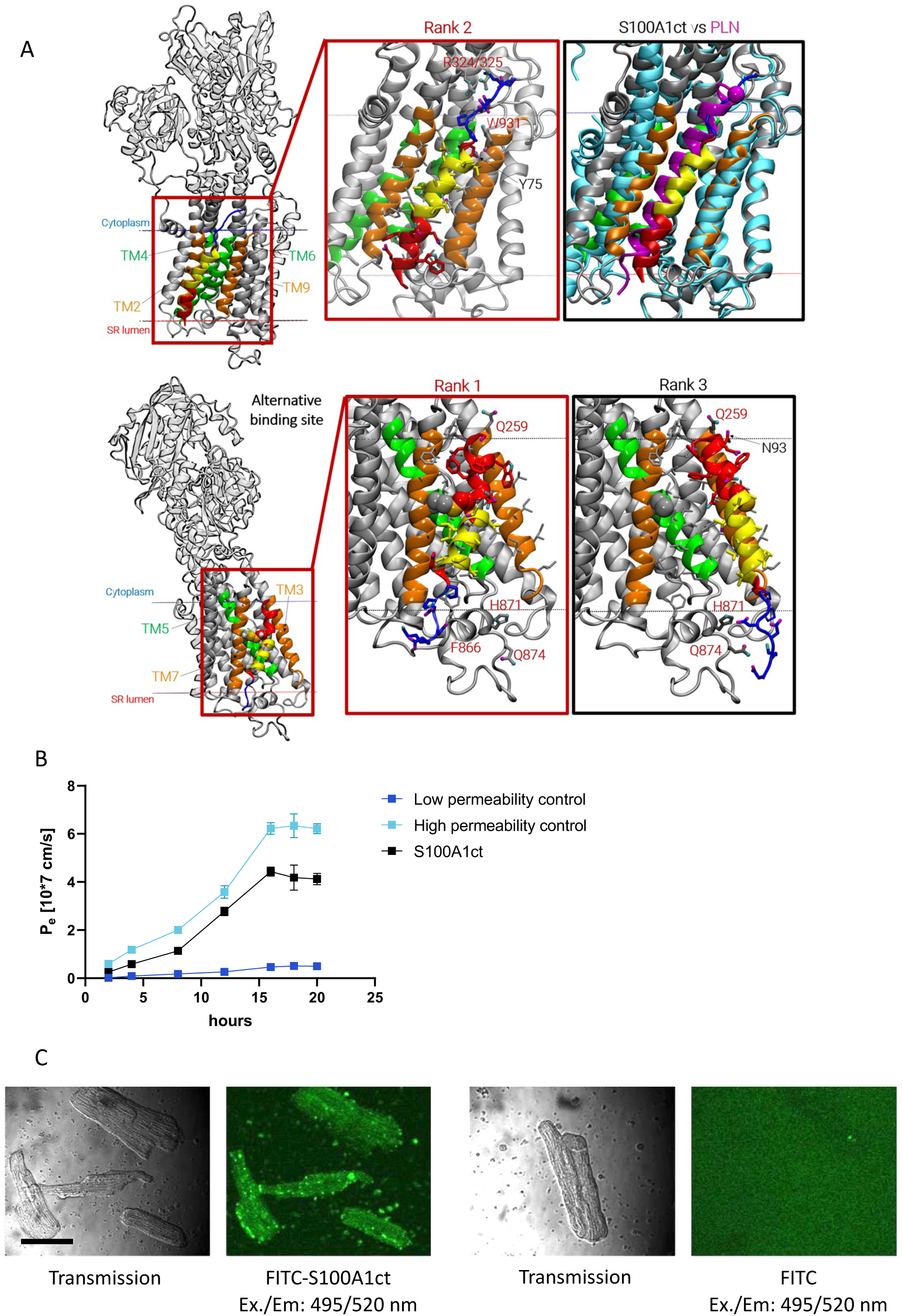
A, Putative structures of complexes of SERCA2a and S100A1ct obtained by computational docking. The three best-ranked docking modes to the SERCA2a E1 state are shown. Corresponding poses for the E2 state can be found in the Supplementary Information. SERCA2a and tagged S100A1ct are shown in cartoon representation. SERCA2a is colored grey, with helices lining the S100A1ct binding sites highlighted in orange or green (green helices are in direct contact with calcium ions). The hydrophilic tag is shown in blue, the apolar sequence stretch of S100A1ct in yellow and the remaining peptide in red. Additionally, the S100A1ct residues are shown as sticks with atom type coloring (carbon atoms: blue/yellow/red, oxygen atoms: magenta, nitrogen atoms: cyan, hydrogen atoms are not shown), with the cysteine residue in the S100A1ct highlighted by van der Waals spheres (sulfur atom: dark yellow). The membrane bilayer is indicated by dashed lines. The cytoplasmic and the luminal sides of the membrane are indicated in blue and red, respectively. Upper row, S100A1ct docked to the PLN binding site (lined by TM2/TM6/TM9; labeled) of SERCA2a. This docking orientation of S100A1ct corresponds to the FMAP 2.0 prediction for the S100A1ct peptide alone in the membrane (see Supplementary Figure 3A). The red inset in the upper row shows residue level interactions of S100A1ct (in case labeled: SERCA2a residues – red labels, S100A1ct residues – black labels). Apolar residues lining the PLN binding side (selected base on visual inspection of the refined docking pose) are shown as thin gray sticks (carbon atoms: gray). SERCA2a residues potentially forming polar contacts (as predicted by PyMOL) in this docking mode with the tag or Y75 of S100A1ct are shown in atom type-colored stick representation. The black inset compares the obtained S100A1ct docking mode with the SERCA2a transmembrane domain (as obtained from https://opm.phar.umich.edu/ for 6JJU)-superposed binding mode of the rabbit SERCA1a (cyan cartoon)/dog PLN (magenta) complex from the PDB-ID 4Y3U. Lower row, Alternative docking mode of S100A1ct to SERCA2a, positioned on the opposite side of SERCA2a to the PLN binding site (i.e., TM3/TM5/TM7). The residue-level representations are identical as in the upper row. Apolar residues lining the docking site are again highlighted. The S100A1ct and SERCA2a cysteine residues C86 and C841, respectively, are shown as either atom type-colored (with or without white hydrogens) or gray spheres. Red inset: rank 1 with disulfide bridge and a potential polar contact of SERCA2a’s F866 to the tag of S100A1ct (other SERCA2a residues are highlighted for comparison to rank 3). Black inset: rank 3 without disulfide bridge but potential polar contacts of SERCA2a residues to the tag and N93 of S100A1ct. B, Parallel artificial membrane permeability assay showing the ability of S100A1ct (1 μM) to cross the 4% lecithin artificial membrane over a time period of 20 hours in comparison to a low and high permeability control. Measurements were carried out in triplicates. Data are given as mean±SEM for P_e_ 10^-7^ cm/s. C, Representative confocal microscopic images of isolated adult ventricular cardiomyocytes subjected to FITC-coupled S100A1ct (1 μM) (left; transmission and fluorescent image) and fluorescent dye only (right; transmission and fluorescent image) for 20 minutes show intracellular accumulation of S100A1ct in cardiomyocytes. Black bar = 50 μm.

### Biological impact of the cell-penetrating S100A1ct peptide on normal cardiomyocyte performance

Given the apparent transmembrane tendency of S100A1ct in our model, we conducted a biomimetic parallel artificial membrane permeability (PAMP) assay to assess the passive permeability characteristics of S100A1ct across a lecithin-based biological membrane. The results, shown in Figure 2B, indicate S100A1ct’s ability to passively cross the biological membrane with a permeability index close to the high permeability control agent. Consistent with our PAMP assay, confocal microscopic imaging showed intracellular uptake of fluorescence-labeled S100A1ct peptide in cultured intact left ventricular (LV) adult ventricular cardiomyocytes (AVCMs) (Figure 2C) that were isolated from healthy rat hearts. In contrast, recombinant human S100A1 (rhS100A1) protein was unable to permeate through the biological membrane and was not taken up by the cardiomyocytes (data not shown). In light of our molecular assays results, these observations posed the question whether S100A1ct peptide might have the ability to impact the contractile function of normal rat AVCMs. Epifluorescent and confocal microscopic intracellular Ca^2+^ imaging in electrically paced and quiescent cardiomyocytes and well as video-edge detection protocols were utilized to determine the impact of S100A1ct on cellular and SR Ca^2+^ handling, Ca^2+^-induced pro-arrhythmic events and contractile performance and to establish dose-efficacy and time-efficacy relationships. Figure 3A describes the dose-dependent enhancement of the Ca^2+^ transient amplitude by S100A1ct in electrical field-stimulated and Fura-2AM loaded normal AVCMs. The increment in cytosolic Ca^2+^ cycling across all groups started after a 3-5-minute incubation period and the *in vitro* dose-efficacy relationship yielded an EC50% concentration for S100A1ct of approximately 120 nM. Neither S100A4ct nor S100Bct peptide altered any of the tested performance parameters (data not shown) which corroborates the specificity of S100A1ct’s *in vitro* actions. The effect of S100A1ct on Ca^2+^ cycling caused an increase of the basal cardiomyocyte twitch amplitude, which was preserved in response to ß-adrenergic receptor (ß-AR) stimulation with isoproterenol (Figure 3B). Subsequent biochemical analyses excluded that S100A1ct causes any phospho-specific post-translational modifications of either phospholamban (PLB) or troponin I (TnI) to generate its performance enhancing effect (Figure S6A and S6B). Furthermore, in line with our biomimetic PAMPA assay results, incubation of normal AVCMs with rhS100A1 protein did not exert any action on cardiomyocyte performance (data not shown). Figure 3C demonstrates that S100A1ct peptide enhanced the SR Ca^2+^ content, which is the major source of cytosolic Ca^2+^ cycling. To next determine whether S100A1ct might convey anti-arrhythmogenic actions, as has also been reported for native S100A1 protein,^31^ we assessed the peptide’s impact on diastolic SR Ca^2+^ leak in AVCMs. Figure 3D depicts the decrease in Ca^2+^ spark frequency by S100A1ct peptide under baseline conditions in quiescent normal AVCMs. Figure 3E shows prevention of diastolic after-contractions by S100A1ct peptide in paced normal AVCMs subjected to a previously described pro-arrhythmogenic isoproterenol/caffeine treatment.^31^ Consistently, Figure 3F illustrates the ability of S100A1ct to abrogate the isoproterenol/caffeine-induced SR Ca^2+^ leak in quiescent rat normal AVCMs, as previously reported for viral-based S100A1 protein overexpression.^31^

**Figure 3.**
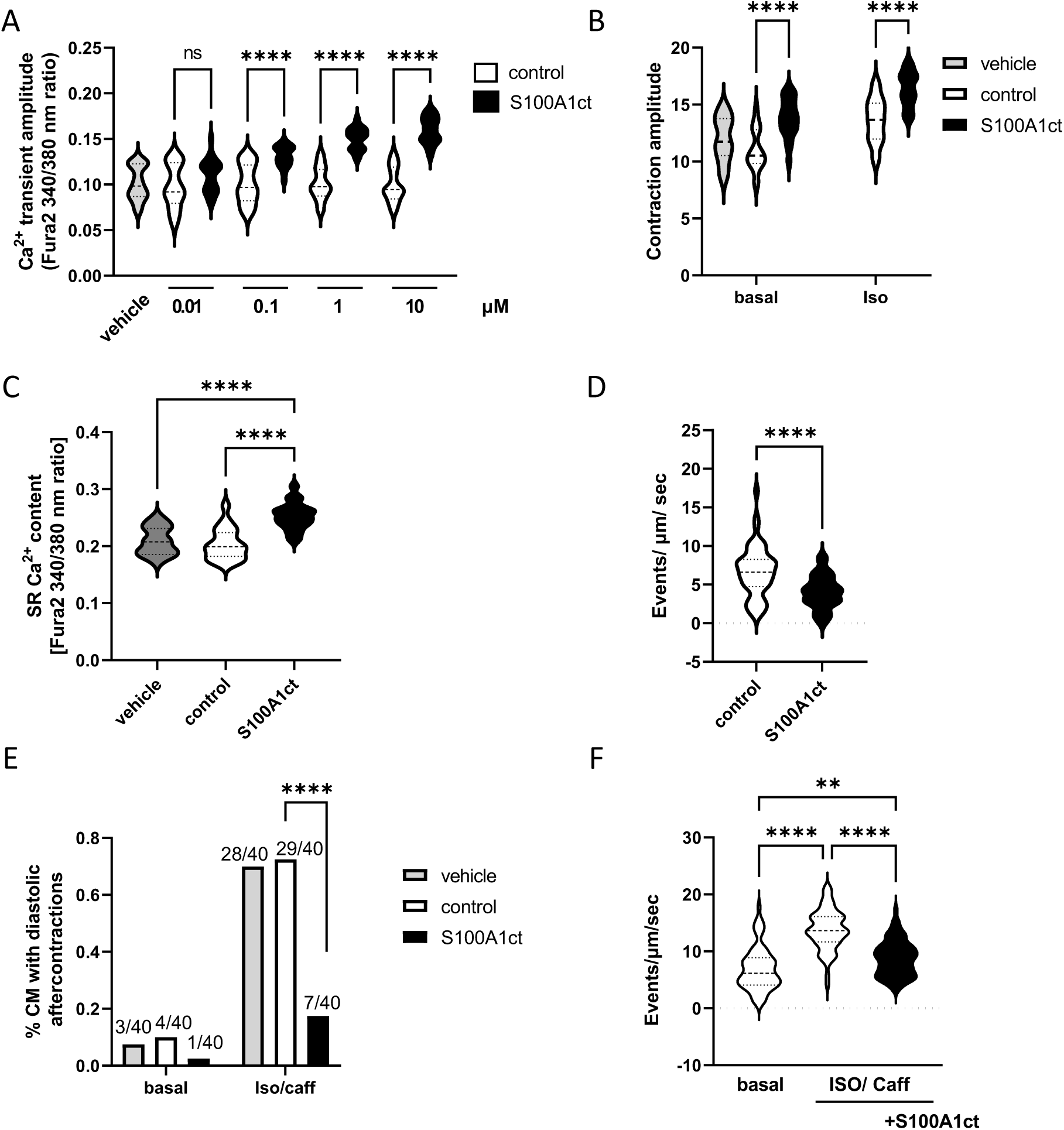
A, Dose-dependent enhancement of the Ca^2+^ transient amplitude in 2 Hz electrically-stimulated and FURA2-AM loaded normal rat LV AVCMs by S100A1ct treatment (0.01, 0.1, 1, 10 μM) versus control. B, Preserved gain-in-function of the contraction amplitude in rat AVCMs by 1 μM S100A1ct in the presence of isoproterenol (0.1 μM) at 2 Hz stimulation. C, Augmented SR Ca^2+^ load due to 1 μM S100A1ct treatment versus control in rat AVCMs, as assessed by the caffeine-pulse protocol. D, Decreased rate of Ca^2+^ spark frequency in quiescent normal rat AVCMs in response to 0,1 μM S100A1ct compared to control. E, Diminished rate of diastolic aftercontractions by 1 μM S100A1ct in 2 Hz electrically stimulated rat AVCMs at basal conditions and in response to a pro-arrhythmogenic isoproterenol/caffeine treatment protocol. F, Abrogated isoproterenol/caffeine-induced SR Ca^2+^ leak by 1μM S100A1ct in quiescent rat AVCMs, as assessed by Ca^2+^ spark frequency rate. For Ca^2+^ transient, SR Ca^2+^ load, diastolic after-contraction and SR Ca^2+^ sparks, n=20, n=20, n=40 and n=30 cells per group, respectively, were used from three different rat heart AVCM isolations. Data are given as mean±SEM. **P=0.0095; ****P<0.001. Vehicle vs. control groups; P=n.s. Control: scrambled peptide, vehicle: aqueous carrier used for scrambled and S100A1ct peptide dilution.

### Therapeutic effect of S100A1ct on failing cardiomyocyte performance and impact of a cardiomyocyte-targeting peptide tag

We next advanced to assessing the therapeutic effect of S100A1ct on failing LV AVCMs isolated from a post-myocardial infarction (post-MI) rat HFpEF model and explanted failing human hearts, as described previously.^21, 22^ Figure 4A and 4B describe the dose-dependent enhancement of the Ca^2+^ transient amplitude by S100A1ct in electrical field-stimulated and Fura-2AM loaded rat and human failing LV AVCMs with EC50% concentrations of 244 nm and 800 nM, respectively. Measurement of SR Ca^2+^ content as published previously^31^ demonstrated that S100A1ct was able of improving this parameter both in rat and human failing AVCMs (Figure 4C 4D). Finally, we tested the potential anti-arrhythmogenic effect of S100A1ct with the previously utilized isoproterenol/caffeine treatment^31^ and showed a significant decrease in the rate of diastolic aftercontractions in rat and human failing AVCMs by S100A1ct (Figure 4E and 4F). With the aim of enhancing the specificity of cardiac S100A1ct actions *in vitro* and *in vivo,* we incorporated a previously published synthetic cardiomyocyte-targeting peptide (CTP) that facilitates cardiomyocyte transduction^41, 42^ into S100A1ct. Figure 5A depicts the sequence of the CTP-S100A1ct (cor-S100A1ct) and CTP-control peptide (cor-scrambled peptide). Of note, cor-S100A1ct displayed similar performance enhancing actions to S100A1ct on Ca^2+^ transient amplitudes both in normal and failing rat AVCMs, respectively, while the control peptide had no effect on cardiomyocyte performance (Figure 5B and 5C). Direct comparison of the dose-dependent impact of both peptides on normal and failing rat AVCMs Ca^2+^ cycling showed approximately 7-fold lower EC50 concentrations cor-S100A1ct (normal AVCMs; 121 nM S100A1ct vs. 18 nM cor-S100A1ct, failing rat AVCMs 244 nM S100A1ct vs. 37 nM cor-S100A1ct) indicating enhanced *in vitro* efficacy for the cardiomyocyte-targeted peptide. In addition, cor-S100A1ct apparently improved the of SR Ca^2+^ content both in normal and failing rat AVCMs (Figure 5D and 5E). We therefore advanced both peptides to small animal models and determined the *in vivo* impact of cor-S100A1ct and S100A1ct on cardiac performance and clinically relevant traits first in healthy mice and then in a previously published post-MI HFrEF mouse model.^35^

**Figure 4.**
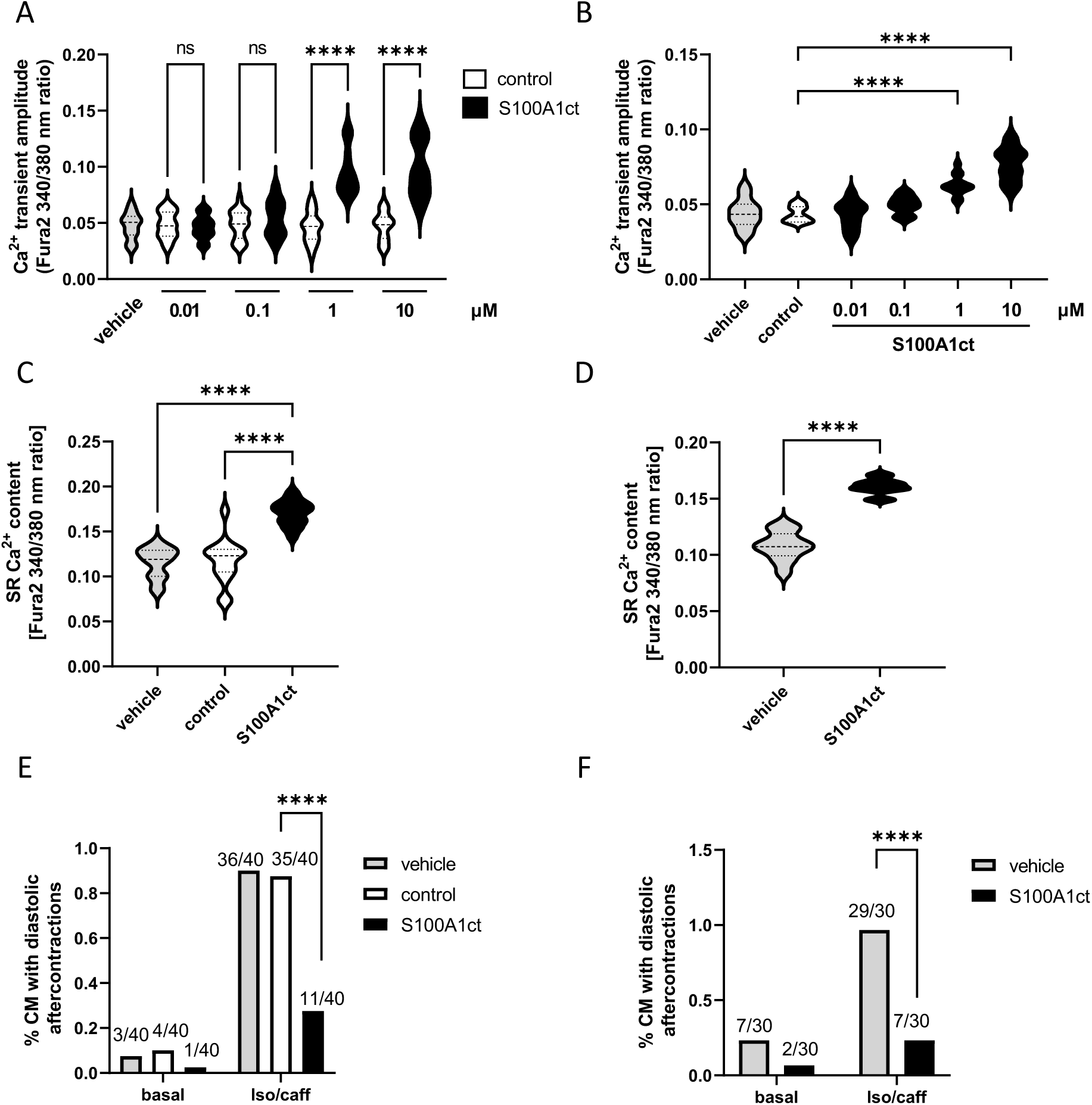
A and B, Dose-dependent enhancement of the Ca^2+^ transient amplitude in failing rat (A) and human (B) LV AVCMs by S100A1ct (0.01, 0.1, 1, 10 μM) treatment versus control. As the control peptide had no effect on failing rat AVCMs at any concentration, its use was limited to a 1 μM control group in the respective human failing AVCM experiment. Measurements in each dose group were performed under steady-state 2 Hz electrical stimulation after a 30 min incubation period. C and D, Effect of 1 μM S100A1ct on the SR Ca^2+^ load both in failing rat (C) and human (D) LV rat AVCMs subjected to the caffeine-pulse protocol. E and F, Decreased rate of diastolic after-contractions by 1 μM S100A1ct both in 2 Hz electrically stimulated failing rat (E) and human (F) LV rat AVCMs subjected to a pro-arrhythmogenic pharmacological isoproterenol/caffeine protocol. As the control peptide showed no difference to vehicle in any experiment of our study, we compared the impact of S100A1ct on SR Ca^2+^ load and after-contractions in failing human AVCMs to vehicle only. For the Ca^2+^ transient, SR Ca^2+^ load and diastolic after-contractions measurements in failing rat AVCMs, n=20, n=15 and n=15 cells per group, respectively, were used from isolations of three post-MI rat hearts. Accordingly, for human failing AVCMs, n=12, n=12 and n=30 cells per group, respectively, were used from isolations from nine explanted human failing hearts. Data are given as mean±SEM. ****P<0.001. Vehicle vs. control groups; P=n.s. Control: scrambled peptide, vehicle: aqueous carrier used for scrambled and S100A1ct peptide dilution.

**Figure 5.**
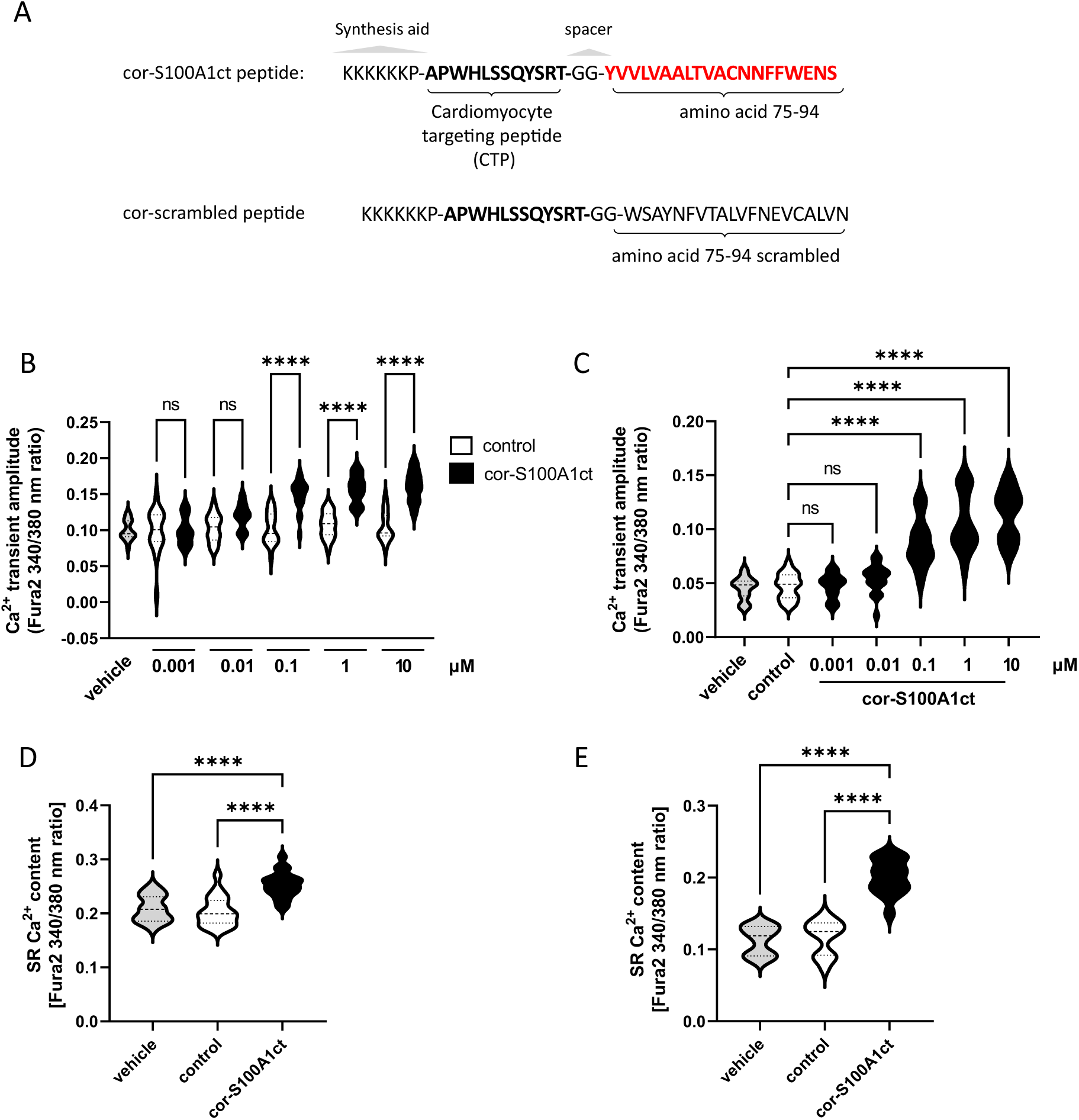
A, Primary sequence of cor-S100A1ct peptide (CTP, cardiomyocyte targeting peptide coupled to S100A1ct) and cor-scrambled peptide (CTP coupled to scrambled peptide). A N-terminal synthesis aid (KKKKKP) and a spacer (GG) were required for scalable synthesis and solubility of both peptides. B and C, Dose-dependent increase of the Ca^2+^ transient amplitude in normal (B) and failing (C) LV rat AVCMs by cor-S100A1ct (0.001, 0.01, 0.1, 1, 10 μM) treatment versus control. Measurements were performed under steady-state 2 Hz electrical stimulation after a 30 min incubation period. As the control peptide had no effect on normal rat AVCMs at any concentration, its use was limited to a 1 μM control group in the respective rat failing AVCM experiment. n=20 cells per normal and failing AVCM group pooled either from 4 different isolations of normal or post-MI rat hearts, respectively. D and E, Impact of 1 μM cor-S100A1ct on the SR Ca^2+^ load both in normal (D) and failing (E) LV rat AVCMs compared to control subjected to the caffeine-pulse protocol. n=20 cells per normal and failing AVCM group pooled either from 4 different isolations of normal or post-MI rat hearts, respectively. Data are given as mean±SEM. ****P<0.001. Vehicle vs. control groups; P=n.s. Control: cor-scrambled peptide, vehicle: aqueous carrier used for cor-scrambled and cor-S100A1ct peptide dilution.

### Impact of S100A1ct-based peptides on cardiac performance in healthy mice

Towards a first translation into a small animal model, we made use of healthy mice, with a study design shown in Figure 6A, to characterize an *in vivo* single dose-efficacy relationship both for the impact of cor-S100A1ct and S100A1ct on cardiac performance. In addition, either the ß-AR blocker metoprolol, the ß-AR agonist isoproterenol or a combination of isoproterenol and caffeine were co-administered to capture the impact of clinically used negative and positive inotropes, respectively. LV catheterization was performed to characterize cardiac contractile performance of S100A1ct-treated hearts. As described previously, mice were instrumented with a pressure-tip catheter inserted into the LV through the carotid artery^35^ for on-line monitoring of cardiac contractility changes for a duration of two hours in response to an incremental i.v. applied cor-S100A1ct dose range. Fig 6B illustrates the dose-dependent increase of the maximum rate (LV+dp/dt_max._) and minimum rate (LV-dp/dt_min._) (data not shown) of LV pressure change for cor-S100A1ct compared to control. Heart rate (HR) in the cor-S100A1ct intervention groups neither increased during the observation period nor showed any difference to the control group during the observation period (Figure 6C). Neither vehicle nor control cor-scrambled peptide treatment had any impact on LV+dt/dt_max_ or HR (Figure S7). End-diastolic and end-systolic LV pressure were not affected by cor-S100A1ct treatment compared to control (data not shown). S100A1ct yielded a similar *in vivo* gain-in-function to cor-S100A1ct but required an approximately 8-fold higher dosage for an equipotent effect on LV+dt/dt_max._ (data not shown). Informed by our dose range finding studies, we then used the 3 μg/kg BW dose for cor-S100A1ct to study its effect on cardiac performance in mice either in conjunction with a ß-AR blocker or ß-AR agonist. Figure 6D shows that the basal cor-S100A1ct-mediated gain-in-cardiac function was preserved when cardiac performance acutely increased in response an i.v. bolus administration of isoproterenol. Vice versa, cor-S100A1ct retained its ability to augment cardiac contractile performance despite the cardio-depressive action of ß-AR blocker metoprolol pre-treatment, as depicted in Figure 6E. Again, basal HR was not affected by cor-S100A1ct and the peptide neither altered the expected rise nor the decline in the heart’s beating frequency caused by isoproterenol or metoprolol (Figure 6E and 6F). S100A1ct, at a dosage of 17 μg/kg BW, exerted comparable biological actions to cor-S1001ct in the presence of isoproterenol and metoprolol (data not shown) indicating that the CTP-tag also improved the *in vivo* efficacy of the synthetic peptide.

**Figure 6.**
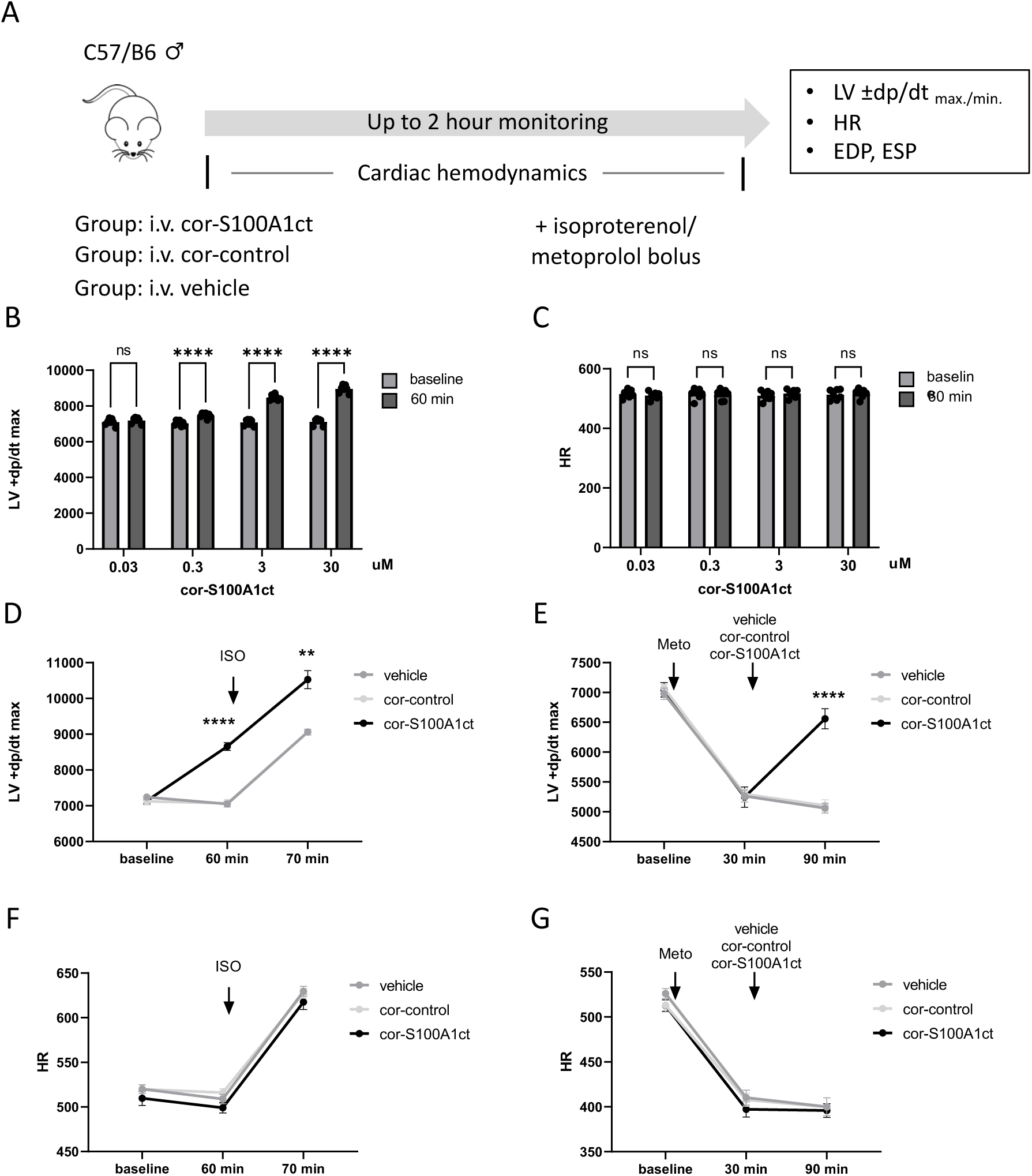
A, Study design to determine the short-term impact of cor-S100A1ct on cardiac performance in male C57/B6 healthy mice. B and C, Impact of a single intravenous (i.v.) dosage range of cor-S100A1ct (0.03, 0.3, 3, 30 μg/kg BW) versus cor-control peptide (0.03, 0.3, 3, 30 μg/kg body weight) on LV+dp/dt_max._ (B) and heart rate (C) 60 minutes after administration. n=7 mice per dosage group. D and E, Preserved gain-in-function in LV+dp/dt_max._ 60 min after a 3 μg/kg BW cor-S100A1ct i.v. dosage compared to cor-control (3 μg/kg BW) in response to a single i.v. dosage of isoproterenol (ISO, 300 pg) (D) or metoprolol (Meto, 90 μg). The effect of ISO in cor-S100A1ct and cor-control treated mice was assessed 10 min after ISO application under steady state conditions. The effect of S100A1ct in Meto-treated mice was measured 60 min after cor-S100A1 and cor-control administration. D and E, the rise and decline in heart rate by isoproterenol or metoprolol was not changed by cor-S100A1ct. n=7 animals per group. Average BW of enrolled healthy mice was 30.66±1.41 g in the control and 30.58±2.03 g in the cor-S100A1ct group. Data are given as mean±SEM. **P=0.025, ****P<0.001. Vehicle vs. cor-control groups; P=n.s. Control: cor-scrambled peptide, vehicle: aqueous carrier used for cor-scrambled and cor-S100A1ct peptide dilution.

### Proof-of-principle for therapeutic activity of S100A1ct-based peptides in post-MI mice with HFrEF

We next advanced cor-S100A1ct and S100A1ct to a post-MI HFrEF model in mice, with the study design depicted Figure 7A, to determine their therapeutic potential *in vivo*. Male and female mice were subjected to experimental MI by ligation of the left anterior descending artery (LAD) as described previously. 48 hours post-MI, surviving mice were either treated with 3 μg/kg BW cor-S100A1ct or 17 μg/kg BW S100A1ct for 12 days by daily intraperitoneal injections and echocardiography in anesthetized mice was performed at day 14 post-MI. Figure 7B demonstrates the decline in LV ejection fraction (EF) in the control group compared to the pre-MI stage that was ameliorated by the 12-day cor-S100A1ct treatment. Moreover, cor-S100A1ct improved post-MI survival in mice (Figure 7C). 14 days post-MI, clinical chemistry biomarker unveiled greater high-sensitive troponin T (hsTnT) levels in control than cor-S100A1ct treated mice (Supplementary Table S3). In light of the antiarrhythmic effects of S100A1ct *in vitro*, we also tested the ability of cor-S100A1ct to mitigate the incidence of tachyarrhythmias and death in response to a previously described pro-arrhythmogenic epinephrine treatment protocol in the 14 day survivors. Figure 7C shows improved survival of post-MI mice with cor-S100A1ct treatment subjected to acute i.v. adrenergic receptor agonist stress (2 mg/kg body weight). S100A1ct, at a dosage of 17 μg/kg BW, exerted comparable therapeutic actions to cor-S1001ct in (data not shown) indicating that the CTP-tag also enhanced the *in vivo* therapeutic activity of the synthetic peptide.

**Figure 7.**
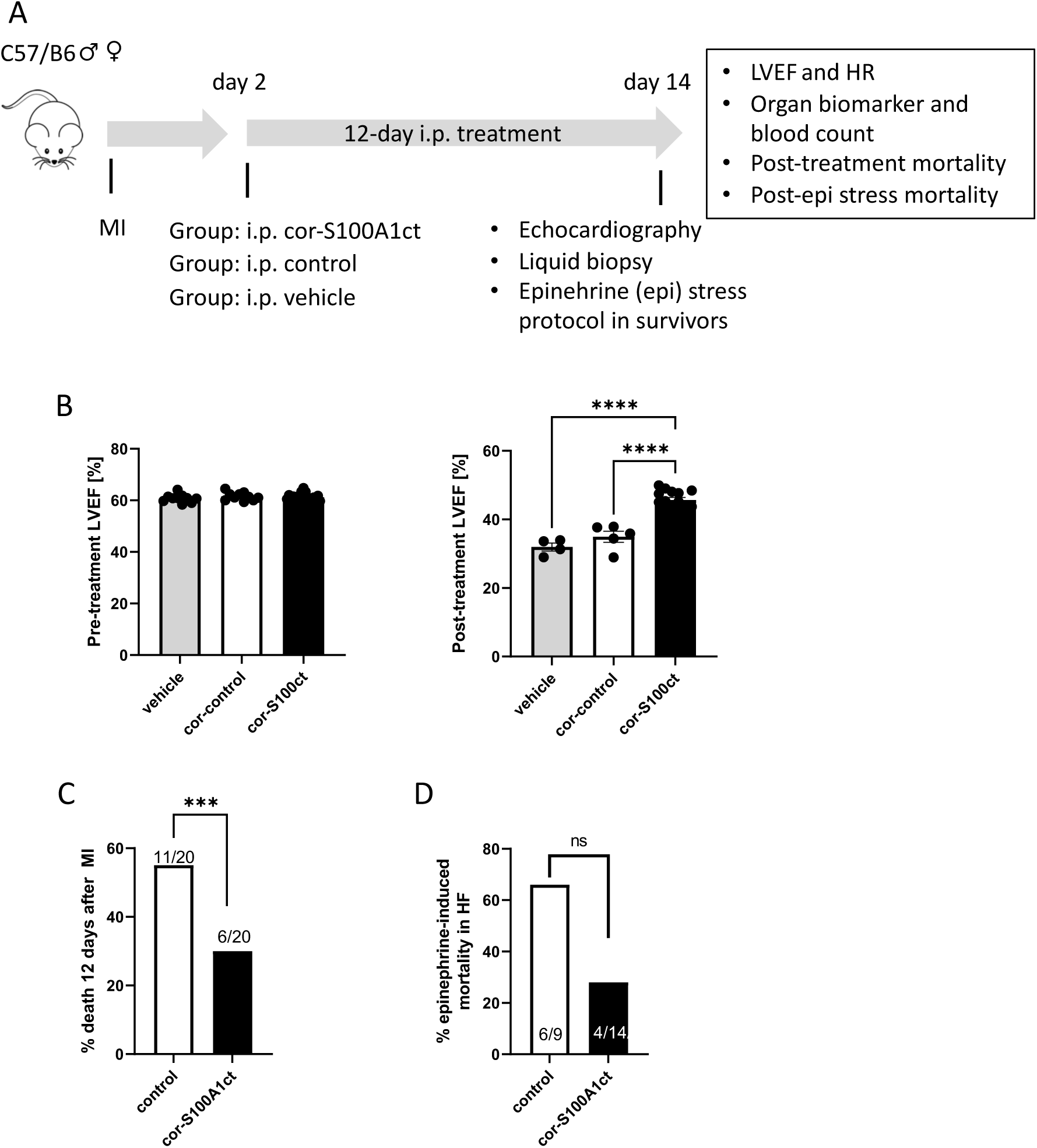
A, Study design to assess the therapeutic impact of cor-S100A1ct on cardiac dysfunction in the post-MI C57/B6 mouse model. 2 days post-MI, surviving animals were treated for 12 days with cor-S100A1ct (3 μg/kg per body weight; BW) (n=20; m/f 10/10) by daily intraperitoneal injections (i.p) and compared to cor-scontrol (n=10; m/f 5/5) and vehicle (n=10; m/f 5/5) treatment. Average BW of enrolled post-MI mice was 30.44±1.01g in the cor-S100A1ct, 30.12±1.82g in the control and 31.32±1.43g in the vehicle group B, Post-treatment impact of cor-S100A1ct on the decline in LVEF in surviving animals (right) compared to the control and vehicle group (vehicle vs. control groups; P=n.s.). LVEF of enrolled animals prior to MI and cor-S100A1ct treatment is shown in the left image. C, Significantly fewer animals had died after the 12-day cor-S100A1ct treatment than in the combined control/vehicle group (control vs. vehicle showed no significant difference and were pooled). D, Acute intravenous epinephrine stress (2 mg/kg body weight) in the 12 days post-treatment showed a trend towards fewer deaths in cor-S100A1ct treated mice than control (combined cor-ontrol/vehicle group). Data are given as mean±SEM. Vehicle vs. cor-control group; P=n.s., ***P=0,012 ****P<0.001. Control: cor-scrambled peptide, vehicle: aqueous carrier used for cor-scrambled and cor-S100A1ct peptide dilution.

### Proof-of-feasibility for therapeutic actions of S100A1ct on cardiac performance in a post-MI pig model

The therapeutic efficacy of S100A1ct-based peptides in a small animal post-MI HFrEF model subsequently prompted a study in a post-MI large animal model to estimate the therapeutic potential of cor-S100A1ct in a model system that more closely approximates human cardiovascular pathophysiology with a study design described in Figure 8A. To enable this study, we used a previously published post-MI cardiac dysfunction model in German farm pigs.^36, 37^ Three weeks after experimental percutaneous catheter-based occlusion of the left circumflex artery (RCA), anesthetized and ventilated animals were either subjected to cor-S100A1ct or control treatment. A single dosage of 150 μg/kg BW cor-S100A1ct was directly infused over a period of approximately 1 minute into the left anterior descending artery (LAD) by a percutaneous microcatheter to minimize the amount of peptide to impact the function of the heart in the human-sized large animal model. Subsequent changes in cardiac performance were assessed over a period of three hours and blood samples were taken prior and after cor-S100A1 treatment to capture potential acute adverse effects of the peptide on organ function by clinical chemistry biomarker and basic blood counts. In addition, an epicardial 12-channel electrocardiogram was recorded to determine the potential influence of cor-S100A1 on critical parameters of the electrical activity of the heart. Figure 8B shows the steady increase in LVSV over time in response to the single dose cor-S100A1ct treatment that reached a significant difference to control after a 3-hour period. Heart rate (HR) was not significantly altered in both groups during the observation period (Figure 8C). We detected a concurrent augmentation in LVEF over time also reaching a significant increase over control 3 hours after administration (Figure 8D). Additional hemodynamic parameters are given in Figure S8 showing, e.g., no difference in end-diastolic filling pressure between groups. Furthermore, the analysis of the epicardial ECG did not yield any abnormalities, e.g., in AV-node transmission, ventricular depolarization or repolarization by the cor-S100A1 treatment, as shown by PQ interval, QRS duration or corrected QT-interval (Figure 8E to 8G). We did not detect any brady- or tachyarrhythmias in either group (data not shown). Also, the assessment of various organ biomarkers by clinical chemistry did not yield any signs of acute cardio- or hepatotoxicity, renal dysfunction or disturbances in glucose and ion homeostasis or non-heparin dependent coagulation parameters by cor-S100A1ct (Supplementary Table S4). In addition, basic blood counts did not show any irregularities in the numbers of leukocytes, erythrocytes or platelets within the 3-hour observation period.

**Figure 8.**
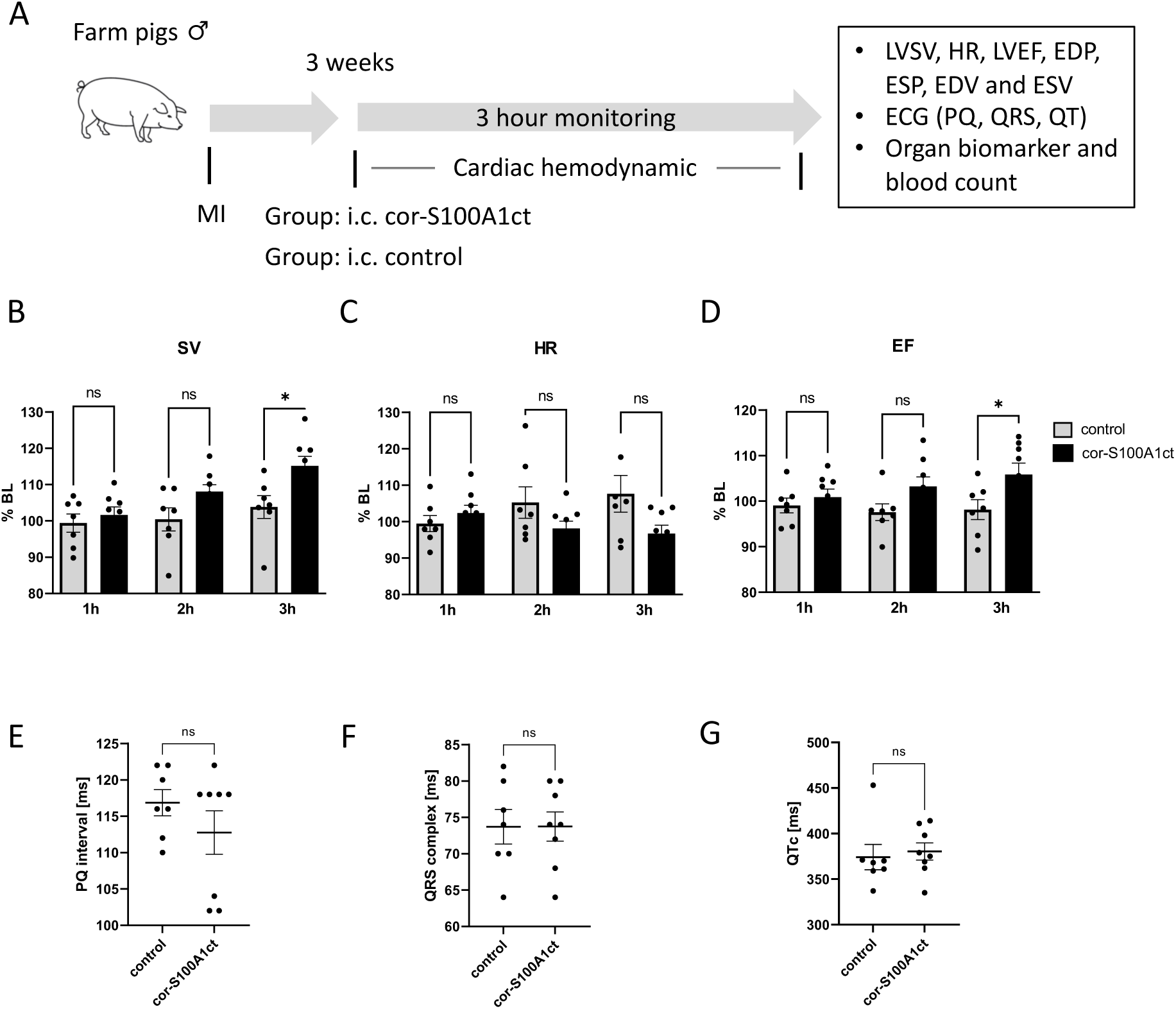
A, Study design to estimate the short-term therapeutic impact of cor-S100A1ct on cardiac dysfunction in a post-MI pig model. 3 weeks post-MI, cardiac function was monitored for 3 hours in anesthetized and ventilated post-MI animals either after a catheter-based intracoronary (i.c) single dosage administration of cor-S100A1ct or control (vehicle). Average body weight (BW) was 40.86±1.82 kg in the control (n=7) and 40±1.37 kg in the cor-S100A1ct (n=8) group. B-D, Impact of 150 μg/kg BW cor-S100A1ct and control treatment on LVSV (B), HR (C) and LFEF (D) over a course of 3 hours, shown as relative hourly changes (%) compared to baseline hemodynamics of each animal. Data are given as mean±SEM as relative (%) change from individual hemodynamic baseline data. LVSV; *P=0.016, LVEF; *P=0.040. E-G, Unchanged cardiac electrical conductivity and activity of porcine hearts after cor-S100A1ct administration compared to control, as assessed by PQ interval (E), QRS duration (F) and corrected QT interval (G). Vehicle: aqueous carrier used for cor-S100A1ct peptide dilution.

## Discussion

Here, we provide evidence for the novel concept that the primary sequence of S100A1’s C-terminal *α*-helix is sufficient to engineer short cell-penetrating peptides, S100A1ct and cor-S100A1ct, that mimic the contractile performance enhancing biological effects of the S100A1 EF-hand Ca^2+^ sensor in cardiomyocytes. As injectable peptides therapeutic with inotropic and antiarrhythmic properties, they proved effective to treat and to restore contractile function of clinically relevant *in vitro* and *in vivo* HFrEF models. By way of comparison with previously published data for S100A1, our results indicate that S100A1ct can mimic the reported molecular actions of experimentally elevated levels of the parent protein in cardiomyocytes.^4, 7^ On the one hand, this conclusion is supported by data from our molecular studies that expand the number of S100A1ct target proteins by SERCA2a and F1-ATPse, as previously demonstrated for S100A1 protein.^17, 21, 23, 24^ The specificity of this finding was corroborated by the inefficacy both of the synthetic S100Bct and S100A14ct peptide and the scrambled control peptide as residues of the C-terminal *α*-helical fragment have the greatest diversity among the S100 protein family members.^5^ S100B and S100A4 were chosen because of an enhanced expression of both proteins in failing myocardium, which is associated with anti-hypertrophic ^43^ and anti-apoptotic^44^ actions, respectively. On the other hand, an equivalent functional impact by S100A1ct seems to occur within a largely intact cardiomyocyte microarchitecture as the peptide enhanced SR Ca^2+^ uptake and decreased myofilament Ca^2+^ sensitivity to the same extent as S100A1 protein in the same chemical-permeabilized cellular and multicellular assay systems.^15^ Targets for S100A1ct in the cardiac contractile apparatus have not yet been identified, but first clues may come from proven binding partners for S100A1. One example is cardiac titin whose passive tension is reduced by the interaction of Ca^2+^ activated S100A1 with the extendable PEVK region.^45, 46^ As such, these findings strongly corroborate the notion that S100A1ct is sufficient to mimic the net effect of Ca^2+^ activated S100A1 protein. As S100A1 however requires full Ca^2+^ saturation to become active, and further post-translational redox modifications are necessary to adjust the threshold for Ca^2+^ binding towards diastolic Ca^2+^ concentrations, S100A1ct - as a pharmacologically usable bioactive lead - is constantly active without the need for such an activation process.

Computational modeling, using SERCA2a as an exemplary molecular target, provided possible structures of S100A1ct/SERCA2a complexes and binding sites. The conformational flexibility of SERCA2a and its characteristics as a multi-span integral membrane protein required the development of a customized computational docking pipeline that integrated global rigid-body docking with subsequent semi-flexible local docking refinement in a membrane-like environment followed by a stepwise pose ranking procedure. The accuracy of the integrative docking pipeline was tested by modeling of the phospholamban (PLN)/SERCA and sarcolipin (SLN)/SERCAcomplexes, and the results were successfully validated against published X-ray crystallographic structures. Modeling of the S100A1ct/SERCA2a complex indicated a preferred interaction between the intramembrane localized S100A1ct and the multi-span transmembrane segments of SERCA2a. Beyond doubt, our *in-silico* model requires experimental validation, but it is interesting to note that the modeling approach placed S100A1ct close to the transmembrane inhibitory PLN binding region on SERCA2a. From this, a possible consequence could be a mechanistic model where S100A1ct could perturb or sterically hinder PLN binding at its inhibitory SERCA2a site and thereby enhance the activity of the SR Ca^2+^ pump. This mechanism will be the subject of future experimental studies. The putative intramembrane location of S100A1ct informed experiments employing a biomimetic PAMPA assay, which is widely used as an *in vitro* model to examine passive transmembrane permeation of drug candidates.^47^ In contrast to hrS100A1 protein, S100A1ct crossed the artificial membrane with a rather high permeability index. The intracellular accumulation of fluorescent-labeled S100A1ct peptide in cultured cardiomyocytes provided strong evidence for the cell-penetrating ability of the synthetic peptide.

From these results, we concluded that extracellularly applied S100A1ct may be able to cross the cardiomyocyte outer membrane and subsequently augment contractility by acting, e.g., on RyR2 and SERCA2a besides other targets. In line with this notion, S100A1ct improved Ca^2+^ handling properties and contractile performance of normal rat AVCMs in a dose-dependent manner, while extracellularly applied S100A1 protein did not affect the functional state of cardiomyocytes. The prompt onset of the S100A1ct effect *in vitro* suggests rapid intracellular accumulation and activity at target sites while the sustained action indicates some resistance against instant intracellular proteasomal and/or peptidase-mediated degradation. As such, S100A1ct actions are highly consistent with previously reported data showing augmented Ca^2+^ transient amplitudes in cardiomyocytes with increased S100A1 protein levels either by acute intracellular installation of the rhS100A protein via a patch pipette^48^ or from cardiac-targeted S100A1 overexpression in mice^49^ and viral-based S100A1 gene transfer.^15^ From this, we concluded that S100A1ct, once inside the cardiomyocyte, must have targeted intracellular RyR2 and SERCA2a activity, which was confirmed in S100A1ct-treated cardiomyocytes by an attenuated Ca^2+^ spark frequency and enhanced SR Ca^2+^ load. S100A1ct also exhibited the same antiarrhythmic properties as intracellularly elevated S100A1 protein levels in cardiomyocytes^31^ by mitigating the ß-adrenergic evoked SR Ca^2+^ leak from Ca^2+^ sensitized RyR2s. Of note, this mechanism prevented pharmacologically-induced diastolic aftercontractions in electrically paced cardiomyocytes, which is a surrogate marker for diastolic Ca^2+^ homeostasis imbalances that, in turn, can cause an electrical instability of cardiomyocytes *in vitro* and *in vivo*.^50^ Most importantly, with these actions, S100Act exerted a dose-dependent therapeutic improvement of contractile and Ca^2+^ handling properties of isolated rat as well as human failing ventricular cardiomyocytes. As isolated human failing cardiomyocytes retain the various molecular derangements that account for Ca^2+^ imbalances and contractile dysfunction in our experimental setting,^51^ it is tempting to speculate that the beneficial effects of S100A1ct in this human *in vitro* disease model could be viewed as a predictor for potential therapeutic efficacy in human failing hearts.

The decision to incorporate a previously published short cardiomyocyte-targeting peptide (CTP) from here on was made with the aim of enhancing the specificity of S100A1ct’s biological and therapeutic cardiac actions *in vivo*. CTP had been generated in an elegant *in vivo* peptide phage display as a vector being able to transduce cardiomyocytes *in vitro* and *in vivo* effectively and specifically.^41, 42^ As both peptides apparently share the ability to cross biological membranes, we deemed CTP as a useful N-terminal conjugate to S100A1ct. Subsequent *in vitro* studies supported this argument by demonstrating that a lower dosage of cor-S100A1ct than S100A1ct exerted equivalent performance enhancing effects in normal and failing rat AVCMs. Hence, S100A1ct and cor-S100A1ct were concurrently examined in healthy mice to determine whether S100A1ct’s favorable *in vitro* molecular mechanisms could be translated into relevant *in vivo* hemodynamic actions and whether cor-S100A1 would allow us to exploit this with a lower systemic dose. Importantly, the intravenous application of increasing amounts of cor-S100A1ct and S100A1ct led to a prompt and dose-dependent enhancement of *in vivo* cardiac contractile performance indices that lasted for several hours. This supports the notion that sufficient amounts of the S100A1ct-based peptides can reach the myocardium after a one-time systemic administration, and that the synthetic peptides could also resist instant degradation from proteolytic systems in the bloodstream, at least for the duration of the observation. S100A1ct’s hydrophobicity may favor, e.g., albumin binding as it has been described for various molecules and drugs^52^ but a systematic assessment of both inotropic peptides pharmacokinetic, biodistribution and *in vivo* half-life is beyond the scope of the current study and warrants future detailed examination.

Of note, the *in vivo* gain-in-function by cor-S100A1ct and S100A1ct was also additive to ß-adrenergic stimulation. This again is consistent with our *in vitro* results showing that S100A1ct neither interferes with the ß-adrenergic pathway nor relies on cAMP-dependent signaling, as was previously demonstrated for S100A1 in numerous model systems.^21, 35, 49^ With the additional finding of a preserved effect of cor-S100A1ct and S100A1ct in response to a pharmacological ß-adrenergic blockade, we envision that the inotropic peptides may be applicable, e.g., to acute cardiac decompensation not only when an inevitable use of a cAMP-dependent inotrope insufficiently improve cardiac hemodynamics but also when the presence of a negative inotropic ß-adrenergic blockade may demand an effective inotropic override by an alternative molecular mechanism. In addition, a stand-alone inotropic support may also be favorable in light of the inotropic and antiarrhythmic potency that clearly distinguishes the S100A1ct-based peptides from traditional cAMP-dependent inotropes. Of note, S100A1ct’s molecular mode of action also greatly differs from previously reported endogenous G-protein coupled receptor-dependent peptides, such as relaxin,^53^ urocortin,^54^ apelin^55^ or calcitonin gene-related peptide,^56^ that were proposed for the treatment of cardiac dysfunction and exert *in vivo* inotropic actions through modulation of cardiomyocyte cAMP levels and/or heart rate increases. In light of a more sophisticated perspective on cardiac inotropes,^57^ S100A1ct may foremost qualify as a cAMP-independent positive calcitrope. Whether cor-S100A1ct and S100A1ct, in light of the parent protein’s beneficial impact on cardiac energetics, may also classify as a mitotrope remains to be determined in future studies. But our results nevertheless seem to affirm the conclusion that the S100A1-derived peptides could be used as a systemically applicable agonist of S100A1 to exploit its beneficial impact on the heart with the ability to adjust effect size and limit potential side effects by a variable dosing regimen.

As intended, the *in vivo* S100A1ct hemodynamic effects could be met by lower systemic dosages of cor-S100A1ct. As recently published characteristics indicate a fast and specific targeting of murine hearts *in vivo* by the CTP peptide and its conjugates,^41, 42^ future studies need to compare the cardiac uptake of cor-S100A1ct and S100A1ct in more detail. Given the high *in vivo* efficacy of cor-S100A1, we advanced both peptides to our first treatment study in mice. Of note, the prolonged treatment with cor-S100A1ct as well as S100A1ct not only ameliorated the progressive decline in LVEF but resulted in a greater survival rate in our murine cardiac disease model. Also, the *in vitro* protective actions of the peptides against a ß-adrenergic receptor-induced pro-arrhythmic Ca^2+^ leakage seem to be reflected in an attenuated rate of epinephrine-triggered fatal tachyarrhythmias, although other, yet unknown mechanisms may also play a role. But the therapeutic effects that could indeed be observed so far with the S100A1ct-based peptides closely resemble the previously reported beneficial outcome of viral-based S100A1 gene therapy in post-MI cardiac disease models by others and our group.^21, 31, 36, 58^ Although cardiac gene therapy, especially by using long-term expressing adeno-associated viruses (AAV), provides the opportunity for sustained efficacy after a one-time treatment, the technology does not yet allow post-treatment adjustment of the effect sizes or even its cessation in case of adverse cardiac and/or off-target effects.^59^ In contrast, a peptide-based treatment principally enables variable dosing and discontinuation to achieve the desired effect size and to cease adverse effects, respectively, but also requires multiple dosing or continuous administration due to its pharmacokinetic properties.^60^ With these considerations and published evidence of therapeutic efficacy for AAV-based S100A1 gene therapy in large animals,^36, 37^ we tested cor-S100A1 in a second therapeutic study in a pig model with post-MI cardiac dysfunction. The transition from a small to large animal cardiac disease model that more closely approximates human cardiovascular pathophysiology and size clearly bears the promise of greater clinical relevance but at the same requires careful consideration of the far more complex logistics to conduct such a study responsibly and safely.^61^ To this end, we decided for a cardiac-targeted route of administration of cor-S100A1 peptide to limit the amount of the difficult to produce peptide and keep the anticipated time until the onset of a potential therapeutic effect as short as possible. The peptide was therefore administered by a brief catheter-based intracoronary infusion and the impact of cor-S100A1ct on LV contractile performance was monitored for a 3-hour period. Importantly, a single dosage of cor-S100A1ct significantly improved both LVEF and LVSV in the porcine cardiac disease model, while heart rate remained unaltered. Despite the short observation time, this promising result largely reflects key findings of S100A1ct-based peptide treatment in our murine disease models and underscores the feasibility of using cor-S100A1ct as a potential peptide therapeutic for the treatment of decompensated heart failure. This either in combination with a clinically used inotrope, such as dobutamine, or even as a stand-alone option as the negative impact on patient outcome even by a short-term clinical treatment with a cAMP-dependent inotrope makes any reduction in the dose of catecholamine or even their avoidance reasonable.^57, 62^ Finally, our *in vitro* and *in vivo* exploratory toxicity assessment did not raise concerns about acute adverse effects of the peptides. In support of this notion, the various clinical chemistry organ biomarkers, coagulation parameters and blood count did not yield evidence for acute organ damage or dysfunction in response to the peptide treatment. In addition, clinically relevant cardiac electrical stability parameters obtained from cor-S100A1ct treated pigs did not provide any signs for atrioventricular or intraventricular conduction or repolarization abnormalities.

Overall, our translational study, using an integrative vertical research trajectory with computational and experimental methodologies, provides comprehensive evidence for the synthetic peptide S100A1ct as a novel and effective systemic treatment option against cardiac contractile dysfunction and tachyarrhythmias of diseased hearts. S100A1ct, as the bioactive lead structure derived from S100A1, seems to draw its efficacy as a peptide therapeutic from the pleiotropic molecular mechanisms of its parent protein and cell-permeability from its hydrophobic molecular scaffold. Conjugation with a cardiomyocyte-targeting peptide seems to improve its cardiac efficacy and specificity clearly prompting a systematic assessment of its pharmacokinetic parameters to inform the design of preclinical studies with relevant pharmacodynamic endpoints to bring the S100A1ct-based peptide therapeutic closer to a first-in-human clinical trial.^63^ Further mechanistic studies of its interactions with its putative molecular targets are expected to aid in understanding of its molecular MoA and to lay the foundation for further rational development as a peptide-drug candidate.

## Supporting information

Kehr et al. Supplement S100A1ct manuscript

## Acknowledgement

This study was supported by grants from the Klaus Tschira Foundation/Informatic for Life consortium (subproject 2 to RCW and PM), the German Center for Cardiovascular Research (DZHK; project 81Z0500101 to PM and Translational Research Project 01KC2008A to PM and HK), German Ministry of Education and Research (Preclinical confirmatory studies, 01KC2008A to PM), National Institute of Heart, Lung and Blood Research (NHLBI; RO1 HL92139 to PM, RO1 HL0616190 and P01 HL147841 to WJK).

## Author Contributions

DK, KV, JB and CB performed large animal experiments. MG, LJ and RS performed computational studies. PM, JR, KS, ME, MV, AJ and EG conducted molecular and cellular analyses, and performed small animal experiments. PJM and AR provided human failing myocardium. PM, RCW and WJK designed experiments. PM, RCW and WJK supervised all studies. HK, WJK, PM, RCW and NF provided institutional support and feedback on the manuscript. PM, RCW, JR, MG and DK wrote and edited the manuscript. All authors approved the final version of this paper.

