## Supplementary material for "S100A1ct: a synthetic peptide derived from human S100A1 protein improves cardiac contractile performance and survival in pre-clinical heart failure models": Kehr et al. Supplement S100A1ct manuscript

1. **Supplemental Materials and** Methods……………………………………………page 1-14

2. **Supplemental Figures and** Tables……………………………………………..page 15-30

### 1. Supplemental Materials and Methods

All animal procedures and experiments were carried out according to the 'Guide for the Care and Use of Laboratory Animals' (National Institutes of Health) and were approved by the local Institutional Animal Care and Use Committee of Baden-Württemberg, Germany and Jefferson University, PH, USA. All mice were housed at 22°C with a 12-hour light, 12-hour dark cycle with free access to water and standard chow. Bay cage types are used for housing of pigs. Per pig, a body weight prescribed floor space of at least 0.5 and 0.7 $m^2$ were provided according to directive 2010/63/EU with straw bedding. The environment was enriched by pellet balls, chains and gnawing rods. Humidity and temperature were kept at 50-60 % and 20-24 °C, respectively. Access to water was unlimited and restricted food provided twice/d (SAF130M).

#### Generation of human recombinant S100A1 protein and S100 peptides

Human recombinant S100A1 protein generation: A recombinant human S100A1 cDNA (accession number X58079) was subcloned into the expression vector pHis-TRX-1 containing a thrombin cleavage side. S100A1 was produced as a fusion protein with a histidine tag at the $NH_2$ terminus by an IPTG-driven expression system in *E. coli*. The fusion protein was purified using a HiTrap chelating HP column (Amersham Pharmacia Biotech) according to the manufacturer's protocol. Subsequently, the histidine tag was removed by thrombin cleavage and purified S100A1 was dialyzed against 25 mM Tris, pH 7.45.[1-3]

S100 and control peptide generation: S100A1ct, scrambled peptide, cor-S100A1ct, cor-scrambled peptide, S100A4ct and S100Bct were custom-made by commercial suppliers (Eurogentec, GeneScript and Bachem) and sequences are given in Figure 1A, S1B and 5A. For stock solutions, lyophilized peptides were dissolved in 20% v/v DMSO and 80% v/v HEPES pH 7.0 and stored until use at -80 C.

**Calcium ATPase activity and sarcoplasmic reticulum calcium uptake assay**

Cardiac microsome preparation: Hearts for microsomal vesicle preparations were obtained from terminally anesthetized male homozygous S100A1 knock out (SKO) mice (n=20; ~160 mg wet weight per heart). After removal of the atria, total ventricular myocardium was homogenized in 5-fold g/ml volume of ice-cold 0.3 M sucrose, 5 mM imidazole- HCI, pH 7.4 (homogenization medium), using a tissue blender at maximum speed for two minutes. The homogenate was centrifuged in a JA l0 rotor (Beckman Instruments, Inc., Fullerton, CA; model J-21 centrifuge) for l0 min at 8,000 rpm (7,700 $g_{max.}$). The pellet was rehomogenized as before in another 15 ml of homogenization medium, again centrifuged as before and the supernatant saved. From this supernatant, the sarcoplasmic fraction (SR) containing microsomal pellet was obtained by centrifugation for 90 min at 30,000 rpm (110,000 $g_{max.}$) in a Beckman 35 rotor. The microsomes were resuspended in 2ml homogenization (~1 mg protein/ml) medium, quick-frozen in liquid nitrogen, and stored at -70"C until use.[3]

SR vesicle $Ca^{2+}$-ATPase activity: With a photometer (Beckman DU 640) adjusted at a wavelength of 340 nm, oxidation of NADH in the absence and presence of thapsigargin (10 µM) was assessed at 37°C in SKO SR microsomes by the difference of the total and basal absorbance to specifically determine SERCA2 activity. The reaction was carried out in a cuvette volume of 1 ml (1.5 µg A23187/ml, 100 mM KCL, 5 mM MgCl2, 0.3 M sucrose, 5 mM $Na_2ATP$, 5 mM HEPES pH 7.0, 5 µM oligomycin and 20 µg of microsomal protein/ml using a coupled-enzyme assay containing 8.5 U/ml pyruvate kinase, 12 U/ml lactic dehydrogenase, 400 µM NADH and 2 mM phosphoenolpyruvate) with pCa 6.0 and all experiments were carried out in triplicate. The rate of ATP hydrolysis (nmol ATP/mg SR protein × min) was estimated from the equation: ATP hydrolysis = ΔOD340nm / Δt × ε × L × S (ΔOD is the decrease in absorbance at 340 nm (due to NADH consumption) during the interval Δt (in min), ε is the NADH extinction coefficient ($6.22 \times 10^6$ ml × $mol^{-1}$ ×·$cm^{-1}$), L the cuvette length (in cm) and S the amount of SR protein added to the cuvette (mg/ml).[4]

SR $Ca^{2+}$ uptake activity: SR $Ca^{2+}$ uptake in chemically-permeabilized cardiomyocytes was performed as described elsewhere with some little modifications.[2,5] Briefly, adult rat ventricular cardiomyocytes ($1x10^6$/ml), isolated as described below, were exposed to 0.1mg/ml β-escin at room temperature for 1 min, β-escin was removed by centrifugation and the cells resuspended in a cytosolic mock solution (in mM: 120 KCl, 5 MgATP, 15 CrP, 1 MgCl, 25 HEPES, 20 $K_2$Oxalate, 0.05 $K_2$EGTA, pH 7.0). Cells were placed in a cuvette (1.5ml) and equilibrated with 0.01mM fura-2 (free salt), 20μM oligomycin and 5μM ruthenium red in mock solution by stirring. $Ca^{2+}$ uptake measurements were started after the addition of $Ca^{2+}$ to a total cytosolic $Ca^{2+}$ concentration ($[Ca^{2+}]_i$) within the cuvette by 67 μM and caused a rapid increase of the free $[Ca^{2+}]_i$ to ~ 1 μM. The decay of the fura-2 fluorescence ratio signal at 340/380nm was measured at 30 Hz and digitized at 10Hz for later analysis to determine the time constant (τ) of extra-sarcoplasmic $[Ca^{2+}]$ decline. The relationship between given calcium concentrations and the resulting fluorescence ratios was established with a series of calibration experiments and analyzed according to [6].

**F1-ATPase activity**

The enzymatic activity of the mitochondrial ATP synthase complex was measured using a spectrophotometric assay as described elsewhere with minor modifications.[7,8] Briefly, aliquots of cardiomyocyte homogenates from adult WT and SKO mice were added to EGTA-buffered activity solution (in mM: 60 sucrose, 50 triethanolamine-HCl, 50 KCl, 4 $MgCl_2$, 2 ATP, 1.5 phosphoenolpyruvate, 2 EGTA, 1 KCN, 0.001 thapsigargin, pH 7.4, with KOH) supplemented with 200 μM NADH, 5 U pyruvate kinase, and 5 U lactate dehydrogenase. Thapsigargin was used to block the activity of the $Ca^{2+}$-sensitive sarcoplasmic reticulum $Ca^{2+}$ ATPase. Free $[Ca^{2+}]$ was adjusted to either 0.2 μM or 0.2 mM, determined by the program REACT. The conversion of NADH to $NAD^+$ was followed spectrophotometrically at 340 nm at 37°C for 3 min. Recombinant human S100A1 protein was added 10 min prior to measurement as indicated. Oligomycin (10 μg/ml) was used to block the mitochondrial ATP synthase. Mitochondrial ATP synthase activity was calculated as the difference between total and oligomycin-insensitive conversion of NADH to $NAD^+$.

**Isolation and culture of ventricular cardiomyocytes**

Human failing cardiomyocytes: Human ventricular myocardium was obtained from 23 patients with severe ischemic heart failure undergoing orthotopic cardiac transplantation (mean echocardiographic EF 18±9%). Our protocols were reviewed and approved by the Thomas Jefferson University Institutional Review Board and University of Heidelberg ethics committee. Briefly, coronary arteries of explanted hearts were immediately perfused with cold, $Ca^{2+}$-free Krebs-Henseleit (KH) solution containing (in mmol/L) glucose 12.5, KCl 5.4, lactic acid 1, $MgSO_4$ 1.2, NaCl 130, $NaH_2PO_4$ 1.2, $NaHCO_3$ 25, and sodium pyruvate 2 (pH 7.4, with NaOH). In the laboratory, a small catheter was placed into the lumen of an artery that supplied a non-infarcted free-wall region of the LV. The perfused myocardial segment was cut from the heart and rinsed for 30 minutes with a nonrecirculating KH solution containing 10 mmol/L taurine. The tissue was then perfused for 30 minutes with 200 mL of KH containing 180 U/mL collagenase, 20 mmol/L 2,3-butanedione monoxime (BDM), 20 mmol/L taurine, and 0.05 mmol/L $CaCl_2$. This solution was recirculated. The collagenase-containing solution was washed from the tissue for 10 minutes with 500 mL KH containing 10 mmol/L taurine, 20 mmol/L BDM, and 0.2 mmol/L $CaCl_2$. The cardiac tissue was then removed from the cannula, and only midmyocardial tissue was minced. The resulting cell suspension was filtered, centrifuged (25*g* for 1 minute), and resuspended in a KH solution (200 mL KH, 1% weight/volume BSA, 10 mmol/L taurine, and 0.25 mmol/L $CaCl_2$). All solutions were equilibrated with 95% $O_2$ and 5% $CO_2$. The temperature was kept at 37°C throughout the isolation. Initial yields of rod-shaped cells were between 7% and 50%. Cells were subjected to $Ca^{2+}$ toleration as described and further cultured in BDM-free M199. All experiments were conducted within 30 hours of cell isolation.[9]

Rat normal and failing cardiomyocytes: Ventricular myocytes from adult rat normal and failing hearts were isolated using a collagenase method as previously published.[10,11] Heart failure (HF) in 10-12 weeks old male Sprague-Dawley rats was induced as described previously employing an approved model of left ventricular (LV) cryoinfarction.[11] All animals subjected to cryoinfarction exhibited comparable impaired cardiac performance after 12 weeks as described previously,[11] reflected by a 18-25% and 25-35% decrease in LV +dP/dtmax and LV -dP/dtmin, respectively, and a 45-55% increase in LV end diastolic pressure accompanied by a 20-25% decrease in LVESP. Animals were anesthetized with sodium pentobarbital (50mg/kg, intraperitoneal) and the aorta was rapidly cannulated after the hearts were excised and perfused with a rate of 8 ml/min in a Langendorff apparatus. Hearts were initially perfused with calcium-free AC medium (ACM) (pH 7.2) consisting of (in mM) 5.4 KCL, 3.5 $MgSO_4$, 0.05 pyruvate, 20 $NaHCO_3$, 11 glucose, 20 HEPES, 23.5 glutamate, 4.87 acetate, 10 EDTA, 0.5 phenol red, 15 butanedionemonoxime (BDM), 20 creatinine, 15 creatine phosphate (CrP), 15 taurine and 27 units/ml insulin under continuous equilibrium with 95% $O_2$/5% $CO_2$. After 5 min the perfusion was switched to ACM plus collagenase (0.5 U/ml, type A; Roche diagnostics GmbH, Germany) for 20-30 min. Finally, perfusion was changed to low $Na^+$, high sucrose Tyrode solution containing (in mM) 52.5 NaCl, 4.8 KCl, 1.19 $KH_2PO_4$, 1.2 $MgSO_4$, 11.1 glucose, 145 sucrose, 10 taurine, 10 HEPES, 0.2 $CaCl_2$ for 15 min. Thereafter left ventricles of digested hearts were cut into small pieces and subjected to gentle agitation to allow for dissociation of cells. Consequently, cells were resuspended in ACM without BDM in which 2 mM extracellular calcium ($[Ca^{2+}]_e$) was gradually reintroduced at 25°C. Cardiac myocytes used for contractility and $Ca^{2+}$ measurements were plated with a density of 30.000 cells/$cm^2$ on laminin coated dishes.

**Myofilament calcium-force relationship assay**

Measurements of the pCa-force relationship in Triton-skinned rabbit ventricular trabeculae (average length/width/thickness: 3,500 ± 300/153 ± 30/124 ± 27 μm) were done as described previously.[2,12] Peptide interventions with the indicated concentrations were done at a sarcomeric length adjusted to 2.2 μm set by laser diffraction.

**Biomimetic parallel artificial membrane permeability assay**

The parallel artificial membrane permeability assay was carried out according to the supplier's protocol (BioAssays Systems; PAMPA-096). S100A1ct was used at a concentration of 1 μM and compared to supplied high and low permeability controls in a time series up to 20 hours. Peak absorbance of S100A1ct (280 nm), high (280 nm) and low (270 nm) permeability controls in donor and receiver wells at indicated times (2, 4, 8, 12, 16 and 20 hours) at room temperature across an artificial 4% lecithin membrane were assessed by an absorbance plate reader. Measurements were carried out in triplicate and the permeability index ($P_e$) for each agent was calculated according to the supplier's protocol.

**Computational modeling of S100A1ct/SERCA2a complexes**

A structure of the S100A1ct peptide was extracted from a snapshot from a molecular dynamics simulation.[13] The N-terminal tag was unstructured, and the rest of the peptide was in an α-helical conformation. The extracted peptide conformation was energy minimized by steepest descent using the CHARMM27 force field[14,15] (as parameters from this force field are also used by ClusPro during its minimization step) and the GROMACS simulation package version 5.0.6[16] (convergence criterion: maximum force smaller than 10 kJ $mol^{-1}$ $nm^{-1}$, treatment of nonbonded interactions: reaction field electrostatics with a long range electrostatic (and van der Waals) cut-off of 1.4 nm and a dielectric constant beyond the cut-off of 78). Two models of human SERCA2a used for docking were generated from crystal structures in the apo (no peptide ligand bound) states: E1-2Ca2+-AMPPCP (PDB-ID: 6JJU[17]; "E1 state") and E2•ATP (PDB-ID: 7BT2[18]; "E2 state"). Metal ions and coenzymes/coenzyme analogues were retained, and all other non-protein atoms removed. The structures were immersed in an implicit membrane by Orientations of Proteins in Membranes (OPM).[19] Missing loops and sidechains were inserted using MODELLER 9v23 (preserving all experimentally determined parts of the structure),[20] generating 1000 models and selecting the model with the lowest DOPE score.[21]

Docking was performed via a customized two-step procedure that was validated against experimentally determined structures for other peptides that bind SERCA2a. Briefly, first, the ClusPro 2.0 webserver[22] was used with default settings and the "Balanced" scoring function for global rigid-body docking of the peptide and SERCA2a. Metal ions were retained for global docking wherever possible (cofactors are removed by default by ClusPro). Next, docked poses were post-processed to account for the presence of the membrane using RosettaMP tools by refining and re-ranking the docked structures using MPDock.[23,24] No cofactors were considered during modeling with RosettaMP tools. Before running MPDock, input models were pre-packed and afterwards 1000 decoys were generated by MPDock. Re-ranking of the 1000 models was performed by (i) discarding the 200 highest energy decoy models based on Rosetta total score, (ii) re-ranking the remaining decoys based on Rosetta interface score ("Isc"), and

(iii) peptide configurational clustering of the top-200 decoy models. Potential polar contacts of the complex models were predicted with PyMOL 2.3.0.[25] Visualization of structures was done with VMD or PyMOL.[26]

**Intracellular $Ca^{2+}$ transients, sarcoplasmic reticulum $Ca^{2+}$ load and diastolic $Ca^{2+}$ leak**

Intracellular $Ca^{2+}$ transients: $Ca^{2+}$ transients in Fura2-AM loaded (2 μM for 20 minutes at 37°C followed by 20 minutes incubation to allow for complete de-esterification of the dye) adult normal and failing cardiomyocytes were obtained 6 hours after plating.[9,11] Measurements were carried out using an inverse Olympus microscope (IX70) with a UV filter connected to a monochromator (Polychrome II, T.I.L.L. Photonics GmbH, Germany) as previously described. Cells were electrically stimulated with a biphasic pulse to contract at 37°C at 2 Hz and excited at 340/380 nm. Epifluorescence emission was detected at 510 nm, digitized, and analyzed off-line with T.I.L.L.VISION software (v. 3.3). Baseline data from 10 consecutive steady-state transients were averaged for analysis of transient amplitude.

Sarcoplasmic reticulum $Ca^{2+}$ load: In a subset of experiments, the total sarcoplasmic reticulum (SR) $Ca^{2+}$ content was immediately assessed after termination of $Ca^{2+}$ transient measurements as described previously.[9,11] After 2 min of electrical stimulation (2 Hz), cardiomyocytes were abruptly exposed to $Na^+/Ca^{2+}$ free solution supplemented with caffeine (20 mM). The peak of the caffeine-induced cytosolic $Ca^{2+}$ rise was used as a semiquantitative index of the SR $Ca^{2+}$ load.

Sarcoplasmic reticulum $Ca^{2+}$ leak: To evoke diastolic SR $Ca^{2+}$ leak-induced after-contractions, respectively, cardiomyocytes were superperfused with a low dose caffeine (0.5mM) and Isoproterenol ($10^{-7}$M) combination as recently described. [9,11] Cells were electrically stimulated with a biphasic pulse to contract at 37°C at 1 Hz. The number of cells with diastolic SR $Ca^{2+}$ leak-induced after-contractions was counted and the peak of the previous systolic $Ca^{2+}$ transient was used as an index of the amplitude.

**Calcium spark analysis**

$Ca^{2+}$ sparks in intact normal and failing adult rat cardiomyocytes were monitored using a Leica SP2, (Mannheim, Germany) laser scanning confocal microscope (LSCM) as described previously.[27,28] Transfected cardiomyocytes were cultured for 24h with HEPES-modified medium 199 as described above. Cells were loaded with Rhod-2AM (Invitrogen 10 μM) for 45 min followed by an additional 45 min interval prior to measurements to allow for complete desterification of the dye. Rhod-2 fluorescence in intact myocytes was excited at 543 nm HeNe laser and measured between 550 and 600 nm applying epifluorescence optics of an inverted microscope with a ×63-1.2 NA water-immersion objective lens. Fluorescence was acquired in line-scan mode at 400 Hz line–1; pixel dimension was 0.232 μM (512 pixels scan–1; zoom =2). The scanning laser line was orientated parallel with the long axis and placed approximately equidistant between the outer edge of the cell and the nucleus/nuclei, to ensure the nuclear area was not included in the scan line. Recorded $Ca^{2+}$ sparks were quantified using an automated detection and measurement algorithm adapted from a previously published method. Up to 15 different cells from 6 animals in each group (approximately 75-90 cells per group) were used for $Ca^{2+}$ spark measurements. Isoproterenol ($10^{-7}$M) and caffeine (0.5mM) were superfused as indicated through a gravity-fed perfusion system and $Ca^{2+}$ spark activity was assessed 3 minutes after stimulation.

**Cellular contraction analysis**

For the measurement of contractile parameters cultured cardiomyocytes were superperfused with modified medium 199 (mM: 5 taurine, 5 D-L creatine, 5 D-L carnitine) (flow rate 3 ml/min) at 37°C and equilibrated with 95% $O_2$/5% $CO_2$.[11,29] Only cardiac myocytes parallelly orientated in respect to the electrical field were chosen for measurements with a video-edge detection system (Crescent Electronics). Images were taken with a sample rate of 120 per second.[30] Steady state twitches were analyzed online (Labview 4.01, National Instruments) to obtain fractional shortening (FS%). Isoproterenol ($10^{-7}$M) was superfused trough a gravity-fed perfusion system and contractions were recorded 5 minutes after β-adrenergic stimulation under steady-state conditions.

**Immunoblotting**

Protein expression analysis was carried out as described in elsewhere. Briefly, cultured cells were rinsed in PBS and scraped off the dish in lysis buffer (PBS, pH 7.4, SDS 1%, and 1 mM EGTA/EDTA) containing a mixture of 1% (v/v) phosphatase inhibitors (Sigma; phosphatase inhibitor mixture I/II) and protease inhibitor (1 tablet/10 ml) (Roche Applied Science; Mini Complete EDTA free protease inhibitor). Protein lysates were subjected to electrophoresis (4-20% tris-glycine gradient gels, ANAMED), transferred to a PVDF membrane (Immobilon FL, Millipore) and probed with appropriate sets of primary antibodies to assess levels of Phospho-Troponin I (Cardiac) (Ser23/24) antibody #4004; Badrilla: PLB pSer16 antibody Catalogue No.: A010-12) and respective total protein (Cell signaling: Troponin I Antibody #4002 and Phospholamban (D9W8M) Rabbit mAb #14562) and fluorescent-labeled secondary antibodies (LICOR Odyssey, 1:10000). Proteins were visualized with a LI-COR infrared imager (Odyssey), and quantitative densitometric analysis was performed by applying Odyssey version 1.2 infrared imaging software. Signals were normalized to housekeeping proteins as indicated in the figure legends and did not different between groups.

**Murine post-myocardial infarction heart failure model**

Myocardial infarction was induced by experimental ligation of the left anterior descending (LAD) as described previously.[31] Under general anesthesia with isoflurane (2%), the heart was exposed via a small left thoracotomy and a suture ligation was subsequently placed at the distal 1/2 of the LAD with 6.0 silk suture. Upon ligation, the heart was immediately placed back to the intrathoracic space followed cy closure of the skin suture and manual evacuation of pneumothoraces. Sham intramyocardial injections (saline) and operation, respectively was performed in a same manner except that the LAD was left unligated. After MI, the animals remained in a supervised setting until fully conscious. Early surgery-related death assigned to severe thoracic bleeding during and less than 6 hours after surgery was similar in all groups (10 %) and excluded from final analysis.

**Porcine post-myocardial infarction cardiac dysfunction model**

Post-myocardial infarction cardiac dysfunction in male neutered German farm pigs was induced by percutaneous transluminal temporary occlusion of the left circumflex artery (LCX) as described elsewhere.[32,33] For induction of ischemic heart dysfunction male neutered German farm pigs were anesthetized with an intramuscular injection of ketamine 15 mg/kg body weight (Ketamin 10 %, Betapharm, Augsburg, Germany) and midazolam 1 mg/kg body weight (Hoffmann-La Roche, Grenzach-Wyhlen, Germany). For perioperative analgesia 0.3 mg buprenorphine (Buprenovet multidose 0.3 mg/ml, Bad Homburg vor der Höhe, Germany) were administered through an IV catheter in the ear vein. After intubation anesthesia was maintained by inhalation of 2-3 % sevoflurane (SEVOrane, AbbVie, Wiesbaden, Germany). Under sterile conditions a 7 F catheter introducer sheath was introduced into the right carotid artery and a 6 F guiding catheter (6 F Judkins right, Cordis, Cardinal Health, Dublin, Ireland) was advanced into the ostium of the left coronary artery. Anticoagulation was achieved by i.v. application of 5000 IU per hour of heparin (Heparin-Natrium LEO 25.000 I.E./5 ml, Leo Pharma, Ballerup, Denmark). Myocardial infarction was induced by temporary (2 h) occlusion of the LCX with a percutaneous transluminal coronary angioplasty (PTCA) balloon (Emerge 3.5 x 15 mm, Boston Scientific, Natick, US) which was placed into the very proximal LCX via a 0.014" guiding wire (Balance Middle Weight, Abbott, Santa Clara, US) and inflated at 6 bar. A potential leakage was ruled out by coronary angiography. 150 mg Amiodaron (Cordarex 150 mg/3 ml, Sanofi GmbH, Frankfurt, Germany) were administered intravenously before (bolus) and during (drip infusion) LCX occlusion minimizing the occurrence of fatal arrhythmias. During the procedure animals were monitored via ECG and capnography. For postoperative analgesia 0.4mg/kg Meloxicam were given once a day for at least 3 days either per intramuscular injection or perorally.

**Peptide Application and Hemodynamic Analysis of Cardiac Function in the Pig Model**

Three weeks after MI intracoronary peptide application under pressure/volume (PV) loop monitoring (ADV 500, Transonic, Ithaca, NY, USA) was achieved. Animals were anesthetized with an intramuscular injection of ketamine 15 mg/kg body weight (Ketamin 10 %, Betapharm, Augsburg, Germany) and midazolam 1 mg/kg body weight (Hoffmann-La Roche, Grenzach-Wyhlen, Germany). An IV catheter was placed in the ear vein. After intubation anesthesia was maintained by inhalation of 2-3 % sevoflurane (SEVOrane, AbbVie, Wiesbaden, Germany), 6-50µg/kg/h fentanyl (Fentanyl-Piramal 0.1mg, Piramal Critical Care, Hallbergmoos, Germnay) and 0.5-1.5 mg/kg/h midazolam (Hoffmann-La Roche, Grenzach-Wyhlen, Germany) i.v.. 3-10 ml/kg/h Ringer's solution (Ringer-Infusionslösung B. Braun, MElsungen, Germnay) were administered intravenously as needed and a suprapubic urinary catheter (Cystofix CH 10, 65 cm, 8 cm puncture cannula, B: Braun, Melsungen, Germany) was placed under the guidance of ultrasound. Three catheter introducer sheaths were placed using two 7F for the right and left carotid arteries and an 8F for the right external jugular vein. Anticoagulation was achieved by i.v. application of 5000 IU per hour of heparin (Heparin-Natrium LEO 25.000 I.E./5 ml, Leo Pharma, Ballerup, Denmark). The PV-Loop catheter (FDH-5018B-E345B) was placed through the left coronary artery in the left ventricle. Blood resistivity was measured, infarcted heart type was chosen, approximate stroke volume was calculated and baseline scan was performed as advised.[34] For peptide application a 6 F guiding catheter (6 F Judkins right, Cordis, Cardinal Health, Dublin, Ireland) was placed in the left coronary artery. A microcatheter (ASAHI Corsair Pro, Asahi Intecc, Seto, Japan) was coated with 1 % human serum albumin (Sigma-Aldrich, St. Louis, Missouri, USA) for 10 min at room temperature and then placed via a 0.014" guiding wire (Balance Middle Weight, Abbott, Santa Clara, US) in the proximal LAD. Peptide (150µg/kg) was diluted in 3 ml 0.9 % saline solution 25mM HEPES pH 6 and administered through the microcatheter. Data acquisition was achieved using iox software from emka technologies (emka technologies, Paris, France).

**Echocardiographic and Hemodynamic Analysis of Cardiac Function**

Transthoracic two-dimensional echocardiography (TTE) in lightly anesthetized mice (tribromoethanol/amylene hydrate; Avertin; 2.5% wt/vol, 8µl/g i.p.) with spontaneous respiration was performed with a 12-MHz probe both in sham and infarcted mice as described in detail elsewhere.[35] TTE in in M-mode was carried out in the parasternal short axis before and after (7 and 28 days) surgical procedure to assess LV diameter and subsequently fractional shortening (FS% = [(LVEDD - LVESD)/LVEDD) × 100]). LV catheterization in mice was conducted as described previously.[32,35] A 1.4 French micromanometer-tipped catheter (SPC-320, Millar instruments, Inc.) was inserted into the right carotid artery and then advanced into the LV. Hemodynamic analysis, including heart rate (beats/min$^{-1}$), LVEDP and LV +dp/dtmax and LV +dp/dtmin, were recorded in closed-chest mode. Peptides and controls were applied via a venous catheter or per intraperitoneal injection at indicated dosages diluted in 3 ml 0.9 % saline solution 25mM HEPES pH 6.0. Epinephrine was given as a single intravenous dosage with a concentration of 2 mg/kg body weight.

**Clinical chemistry, blood count and electrocardiogram**

Clinical chemistry organ biomarker and basic blood count from murine and porcine blood, plasma and serum samples was conducted by Thomas Jefferson Hospital laboratory services and Heidelberg University Hospital clinical chemistry unit. For porcine ECG evaluation 2nd lead of 12-channel-ECG (Cardiovit CS-200, Schiller Medizintechnik GmbH, Feldkirchen b. München, Germany) were used. QT times were corrected using Fridericia's formula (QTcF = QT interval/RR 1/3).[33]

### 2. Supplemental Figures

#### Supplementary Figure 1

**S1A**

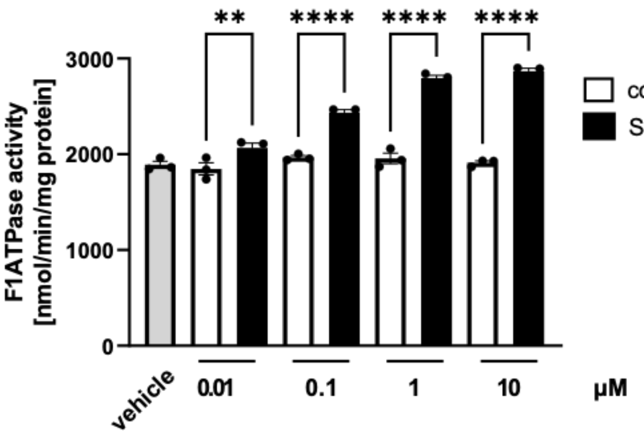

**S1B**

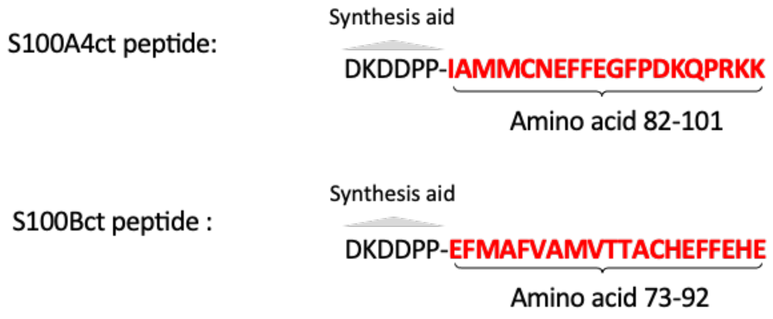

**S1C**

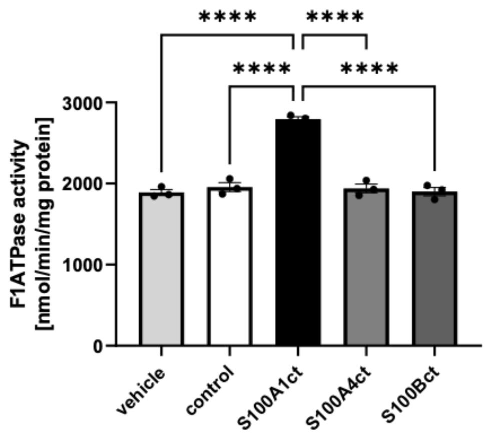

**S1D**

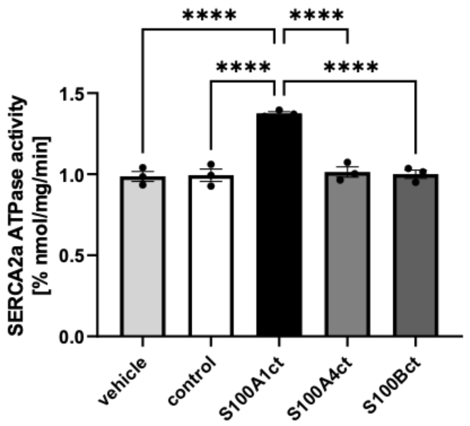

**Supplementary Figure S1 A-D:** A, Dose-dependent augmentation of hydrolytic F1-ATPase activity by S100A1ct peptide. Measurements were carried out in triplicates. Data are given as mean±SEM. *P=0.031; ****P<0.001. B, Primary sequences of the S100A4ct and S100Bct peptides encompassing the respective C-terminal domain of the native proteins. D and E, Neither S100A4ct nor S100Bct altered activity of F1-ATPase or SERCA2a compared to S100A1ct. Measurements were carried out in triplicates. Data are given as mean±SEM. ***P<0.001. Control: scrambled peptide, vehicle: aqueous carrier used for scrambled and S100A1ct peptide dilution.

**Supplementary Figure 2**

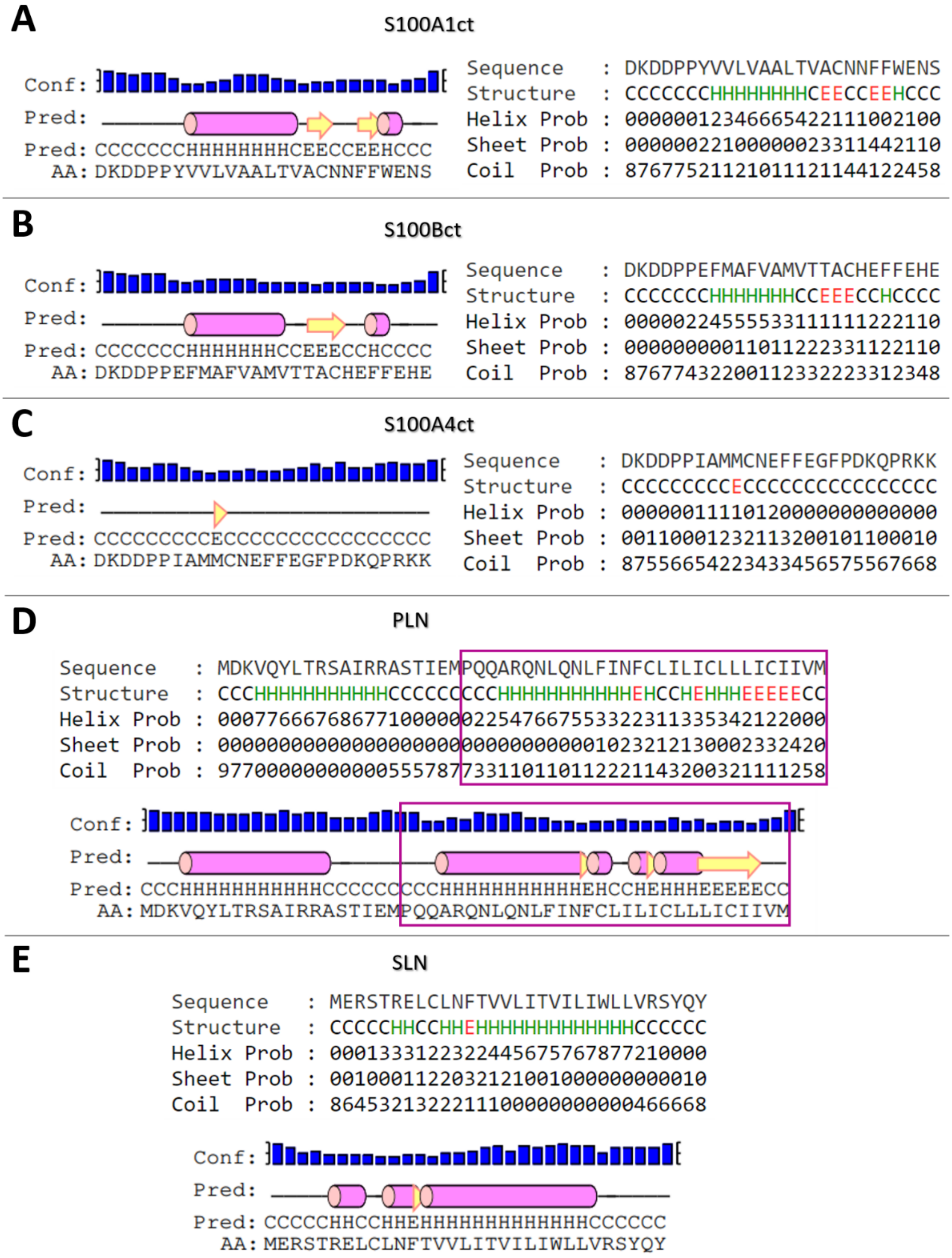

**Supplementary Figure S2 A-E:** Peptide secondary structure prediction results obtained with PEP2D, a secondary structure prediction program specifically tailored for peptides, for A) S100A1ct, B) S100Bct, C) S100A4ct. PLN (D; cardiac phospholamban from dog) and SLN (E; rabbit sarcolipin) are provided as controls. The part of the PLN sequence that encompasses the transmembrane helix of PLN, e.g., in the PLN/SERCA1a structure PDB ID 4Y3U, is highlighted in the magenta box (also see the FMAP 2.0 Server prediction in Supplementary Table S1). For SLN, the sequence provided corresponds to a helix in the structure with PDB ID 4H1W. H: helix, E: sheet, C: coil.

**Supplementary Table S1:** Predictions of propensity for transmembrane helix formation of S100 peptides and peptide controls. The FMAP 2.0 server was used to predict the preferred position of a transmembrane helix in the input sequences, the positioning of the helix in a lipid bilayer (corresponding to an endoplasmatic reticulum (ER) membrane environment), and the corresponding energetic stability and transfer energies at pH 7 and a temperature of 300 K. PLN (phospholamban) and SLN (sarcolipin) were chosen as controls as they both have a well-characterized transmembrane helix. The positioning of the predicted α-helical segments with respect to the lipid bilayer is shown in Supplementary Figure S3. Together with the secondary structure predictions with PEP2D (Supplementary Figure S2), these results show that of the S100 peptides, only S100A1ct is predicted to have an alpha-helix that crosses a membrane bilayer.

| Peptide | Sequence with predicted α-helical segments in bold and cyan | ER membrane binding energy of peptide [kcal/mol] | Stability of helical segment relative to coil in water [kcal/mol] | Transfer energy ($\Delta G_{transfer}$) of helical segment to membrane [kcal/mol] | Tilt angle [°] | Depth/Thick-ness [Å] |
|---|---|---|---|---|---|---|
| **S100A1ct** | DKDDP**PYVVLVAALTVACNNFF**WENS | **-7.8** | **-6.4** | **-9.0** | **7** | **28.2** |
| S100Bct | DKDDPP**EFMAFVAMVTTACHEF**FEHE | -7.7 | -6.5 | -11.1 | 83 | 7.2 |
| S100A4ct | DKDDP**PIAMMCNE**FFEGFPDKQPRKK | -4.8 | -1.4 | -6.0 | 73 | 6.5 |
| Scrambled Peptide | DKDDPPWS**AYNFVTA**LVFNEVCALVN | -6.1 | -1.1 | -5.1 | 83 | 3.8 |
| | DKDDPPWSAYNFVTAL**VFNEVCAL**VN | | -3.1 | -7.3 | 77 | 5.7 |
| PLN$ | PQQARQN**LQNLFINFCLILICLLLICIIV**M | -21.1 | -21.0 | -23.9 | 15 | 30.0 |
| SLN+ | M**ERSTRELCLNFTVVLITVILIWLLVRSY**QY | -22.3 | -22.0 | -25.8 | 24 | 31.1 |

$Representing the C-terminal, transmembrane helix-forming part of cardiac PLN from dog as bound to rabbit SERCA1a in the structure PDB ID 4Y3U.
+Representing SLN from rabbit as bound to rabbit SERCA1a in the structure PDB ID 4H1W.

**Supplementary Table S2:** Results of the computational pipeline applied to docking S100A1ct to SERCA2a. The recommended ranking criterion for ClusPro is the cluster size, which is also used to sort the table. Re-ranking with the MPDock pipeline is indicated with cyan labels for the first seven ranks. Pose features were assigned by manual inspection of ClusPro docking poses (see footnotes). n.d.: no re-ranking with MPDock could be performed due to the lack of a transmembrane span. The very high Rosetta "Isc" scores for ClusPro cluster rank 3 are due to formation of a potential disulfide bridge.

| Clus Pro cluster rank | Pose features ("binding site", "orientation")* | **Clus Pro clust -er size** | ClusPro energies | | Rosetta "Isc" (Interface score) energies after refinement of ClusPro pose with local docking via MPDock | | | |
|---|---|---|---|---|---|---|---|---|
| | | | Lowest energy in cluster | Energy of cluster center | **Lowest energy pose from the biggest conformational cluster obtained with the MPDock pipeline [REU]** | **Rank after refine -ment step** | Average energy of the biggest conform-ational cluster from the MPDock pipeline [REU] | Overall lowest energy from the top-200 poses used for conform-ational clustering [REU] |
| **Docking to the E1 state** | | | | | | | | |
| 0 | TM2/TM6/TM9, ↑ | **173** | -1288.9 | -1118.8 | **-42.81** | ② | -35.93 | -43.71 |
| 1 | TM1/TM2, ↑ | **148** | -1197.2 | -1059.2 | **-27.92** | **15** | -24.71 | -33.91 |
| 2 | TM1/TM2, ↓ | **128** | -1201.8 | -1073.2 | **-26.21** | **17** | -23.29 | -34.42 |
| 3 | TM3/TM5/TM7, ↓ | **76** | -1106.8 | -998.1 | **-60033.46** | ① | -60027.1 | -60036.83 |
| 4 | TM1/TM2, ↓ | **75** | -1167.0 | -1053.0 | **-25.38** | **1** | -21.46 | -35.14 |
| 5 | TM1/TM2, ↑ | **62** | -1207.4 | -1016.6 | **-33.07** | **8** | -28.94 | -40.35 |
| 6 | TM2/TM6/TM9, ↑ | **53** | -1151.3 | -1029.9 | **-27.42** | **16** | -24.72 | -33.90 |
| 7 | TM2/TM6/TM9, ↓ | **44** | -1109.0 | -999.1 | **-37.55** | ⑤ | -33.91 | -39.34 |
| 8 | TM2/TM6/TM9, ↑ | **43** | -1125.2 | -1034.1 | **-35.06** | ⑦ | -31.13 | -39.70 |
| 9 | TM2/TM6/TM9, ↓ | **36** | -1129.4 | -989.0 | **-30.37** | **13** | -25.80 | -35.62 |
| 10 | TM2/TM6/TM9, ↑ | **19** | -1055.9 | -990.9 | **-25.84** | **18** | -23.85 | -38.32 |
| 11 | TM2/TM6/TM9, ↑ | **17** | -1076.5 | -1010.7 | **-37.15** | ⑥ | -32.98 | -38.84 |
| 12 | TM1/TM3/TM4, ~ | **17** | -1079.3 | -1079.3 | **n.d.** | **n.d.** | n.d. | n.d. |
| 13 | TM1/TM2, ↑ | **16** | -1143.5 | -1005.0 | **-30.50** | **12** | -27.33 | -39.55 |
| 14 | TM2/TM6/TM9, ↓ | **15** | -1052.3 | -1036.3 | **-38.32** | ④ | -33.98 | -42.59 |
| 15 | TM2/TM6/TM9, ↓ | **15** | -1066.0 | -1002.1 | **-31.56** | **11** | -27.83 | -39.46 |
| 16 | TM3/TM5/TM7, ↑ | **15** | -1070.8 | -1070.8 | **-31.90** | **10** | -29.31 | -37.03 |
| 17 | TM3/TM5/TM7, ↓ | **13** | -1066.1 | -989.5 | **-40.49** | ③ | -35.36 | -40.49 |
| 18 | TM8/TM9/TM10, ↓ | **10** | -1051.6 | -1051.6 | **-30.18** | **14** | -28.01 | -40.85 |
| 19 | TM1/TM2, ↓ | **6** | -1077.4 | -1013.2 | **n.d.** | **n.d.** | n.d. | n.d. |
| 20 | TM2/TM6/TM9, ↓ | **1** | -995.7 | -995.7 | **-32.70** | **9** | -29.99 | -36.55 |
| **Docking to the E2 state** | | | | | | | | |
| 0 | TM2/TM6/TM9, ↓ | **199** | -1281.6 | -1221.8 | **-39.14** | **9** | -34.73 | -43.31 |
| 1 | TM8/TM9/TM10, ↑ | **100** | -1094.7 | -995.8 | **-29.24** | **13** | -27.44 | -33.31 |
| 2 | TM1/TM3/TM4, ↓ | **98** | -1189.4 | -1189.4 | **-44.65** | ⑤ | -40.59 | -51.75 |
| 3 | TM2/TM6/TM9, ↑ | **90** | -1159.9 | -1013.7 | **-44.81** | ④ | -33.83 | -44.81 |
| 4 | TM2/TM6/TM9, ↑ | **90** | -1163.7 | -1016.0 | **-44.55** | ⑥ | -33.02 | -44.55 |
| 5 | TM2/TM6/TM9, ↑ | **85** | -1162.3 | -1081.9 | **-40.04** | ⑦ | -34.59 | -41.59 |
| 6 | TM3/TM5/TM7, ↓ | **79** | -1226.4 | -994.0 | **-49.46** | ② | -43.24 | -49.46 |
| 7 | TM3/TM5/TM7, ↑ | **68** | -1082.6 | -1082.6 | **-58.40** | ① | -48.29 | -58.40 |
| 8 | TM8/TM9/TM10, ↓ | **42** | -1124.6 | -1009.4 | **-25.72** | **15** | -24.36 | -40.75 |
| 9 | TM2/TM6/TM9, ↓ | **35** | -1110.7 | -1000.2 | **-38.62** | **10** | -34.71 | -38.62 |
| 10 | TM3/TM5/TM7, ↑ | **30** | -1067.7 | -1067.7 | **-33.21** | **12** | -30.38 | -40.48 |
| 11 | In luminal membrane layer adjacent to SERCA2a | **26** | -1145.0 | -1145.0 | **n.d.** | **n.d.** | n.d. | n.d. |
| 12 | TM3/TM5/TM7, ↓ | **15** | -1067.9 | -1037.6 | **-48.80** | ③ | -41.17 | -48.80 |
| 13 | TM8/TM9/TM10, ↑ | **11** | -1071.5 | -990.5 | **-39.67** | **8** | -35.24 | -46.35 |
| 14 | TM2/TM6/TM9, ↓ | **2** | -998.1 | -998.1 | **-37.63** | **11** | -33.41 | -40.30 |
| 15 | TM2/TM6/TM9, ↓ | **2** | -1050.7 | -1011.7 | **-28.96** | **14** | -28.96 | -36.99 |

*Visualizations of the corresponding ClusPro docking poses can be found in Supplementary Figure S4, grouped according to assigned binding/docking sites. TM2/TM6/TM9 denotes the "classical PLN binding site" of SERCA, as found, e.g., in the crystal structure of the dog PLN/rabbit SERCA1a complex with PDB-ID 4Y3U. The other combinations of transmembrane helices denote alternative dockings sites found within the SERCA transmembrane region. Human SERCA2a transmembrane helices are defined as follows: TM1, amino acids (aa) 59-78; TM2, aa 87-107; TM3, aa 259-278; TM4, aa 288-311; TM5, aa 758-780; TM6, aa 788-806; TM7, aa 833-853; TM8, aa 895-915; TM9, aa 929-948; TM10, aa 963-986 (source: https://opm.phar.umich.edu/; PDB-IDs: 6JJU and 7BT2). Docking sites are defined rather “roughly”, as poses can be shifted within as specific docking site. “↑” and “↓” indicate whether the N-terminus points towards the direction of the cytosolic domain of SERCA, i.e., the orientation that is present in the corresponding crystal structures of dog PLN and rabbit SLN with rabbit SERCA1a (i.e., PDB-IDs 4Y3U and 4H1W, respectively) or if the N-terminus points in the opposite direction, i.e., towards the luminal loops of SERCA. “~” indicates that the N- to C-terminal axis of the α-helix of the S100A1ct pose is not oriented perpendicular to the membrane bilayer (as e.g., for PLN binding) but is parallel to the membrane bilayer.

**Supplementary Figure 3**

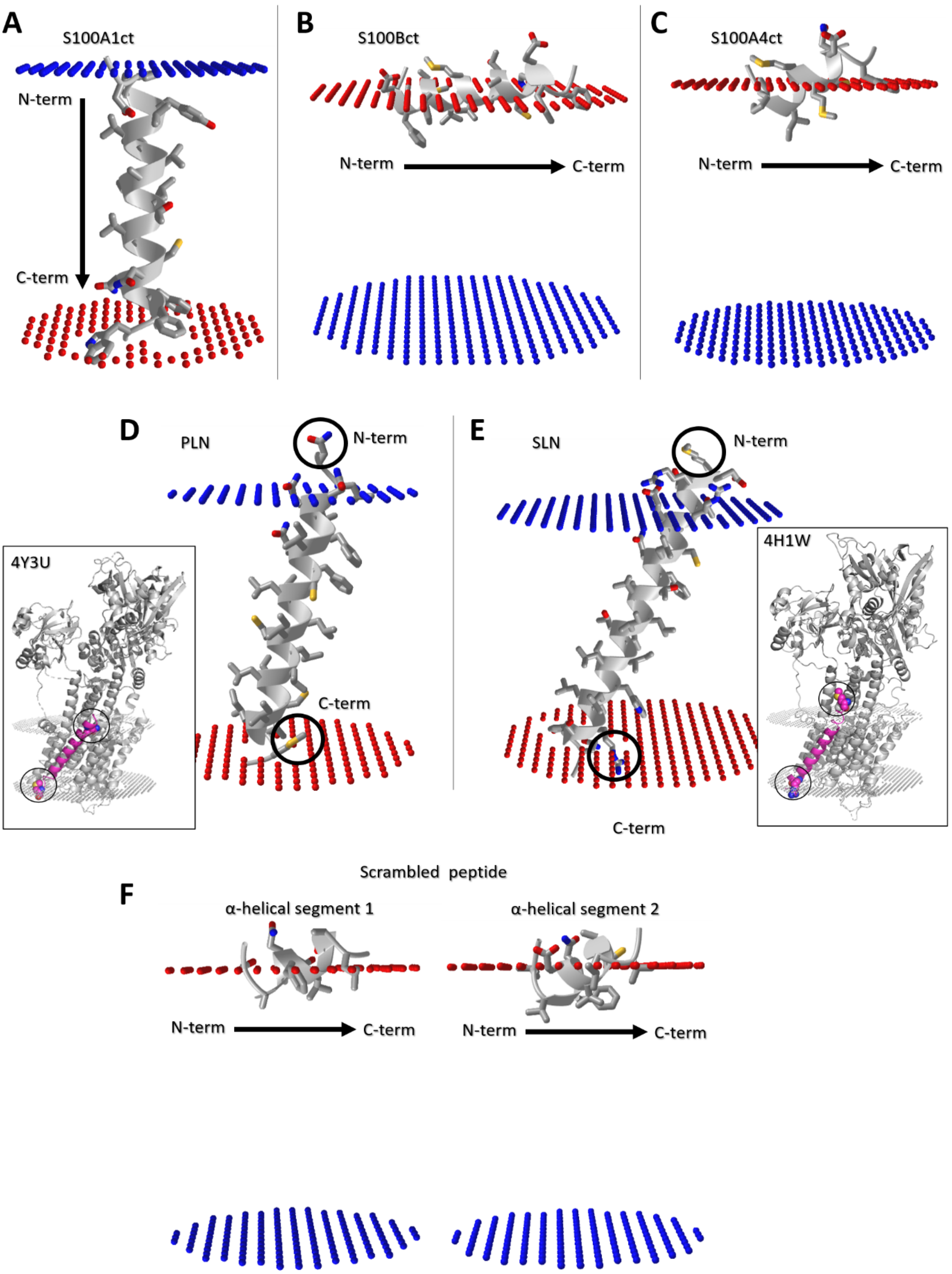

**Supplementary Figure S3 A-F:** Visualization of positioning of the predicted α-helical segments with respect to the lipid bilayer for the FMAP 2.0 server predictions given in Supplementary Table S1. A) S100A1ct, B) S100Bct, C) S100A4ct. PLN (D; cardiac phospholamban from dog) and SLN (E; rabbit sarcolipin) are provided as controls. The predictions for the scrambled peptide are shown in (F). The predicted α-helical segments are depicted in gray cartoon representation with N- and C-termini labelled, and the amino acid residues are shown as sticks in atom type coloring (oxygen atoms in red, nitrogen atoms in blue, sulfur atoms in yellow, and carbon atoms in gray). The lipid bilayer region is indicated between the blue and red dot surfaces. For the PLN and SLN FMAP 2.0 predictions, the crystal structures with the PDB IDs 4Y3U and 4H1W, respectively, showing PLN/SLN in their SERCA1a bound states, are provided as insets for comparison. In the insets, SERCA is shown in gray cartoon and PLN/SLN in magenta cartoon representation and the bilayer is indicated by gray dot surfaces. Residues adjacent to the transmembrane helices are highlighted by black circles in the panels D/E and the corresponding insets, where they are shown in van der Waals sphere representation. The N- and C-terminal orientation of the peptides in the panels D/E and the respective insets is the same. In the inset of panel D, only the part of PLN that is unambiguously resolved in the crystal structure is shown (i.e., ‘PQQARQNLQNLFINFCLILICLLLICIIVM’).

**Supplementary Figure 4**

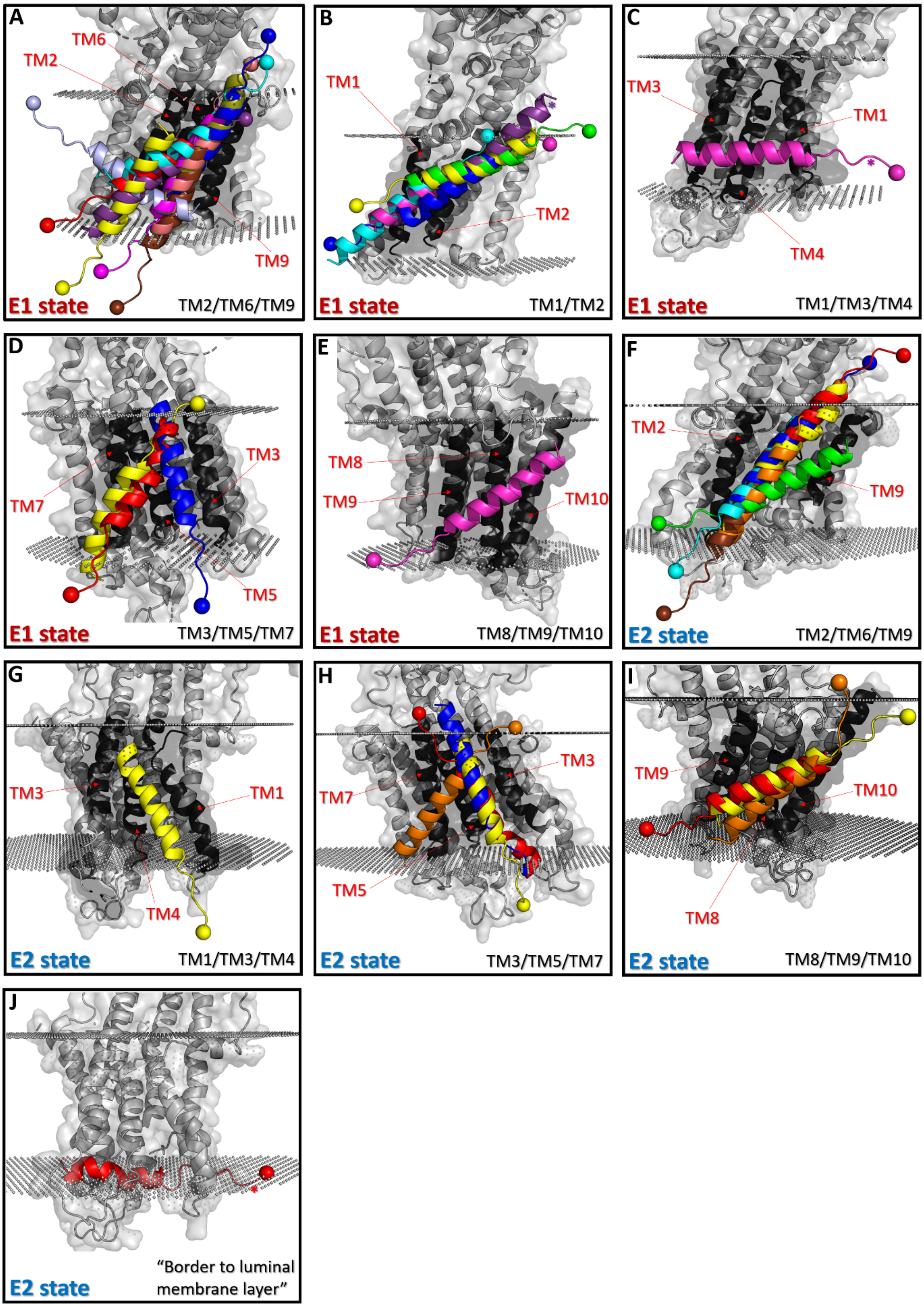

**Supplementary Figure S4 A-J:** The S100A1ct peptide is predicted to dock to the transmembrane domain of SERCA2a by ClusPro. Poses of S100A1ct obtained with ClusPro (shown in differently colored cartoon representations) for docking to the E1 state (i.e., PDB-ID 6JJU) (panels A-E) and the E2 state (i.e., PDB-ID 7BT2) (panels F-J) of human SERCA2a are shown, grouped according to the docking sites listed in Supplementary Table S2. The respective SERCA2a crystal structure is shown in gray cartoon and transparent surface representation (i.e., either 6JJU or 7BT2 with the membrane bilayer position as obtained from https://opm.phar.umich.edu/ indicated by gray dots). Note that the membrane bilayer is only shown for reference and no membrane environment was considered in the dockings performed with ClusPro. SERCA2a transmembrane helices corresponding to respective docking sites are colored in black and labeled in red. The N-terminal Cα atom of the S100A1ct peptide is highlighted by a sphere. ClusPro docking poses that could not be re-ranked with MPDock pipeline are marked with an asterisk. E1 state ClusPro clusters: A) 0 (pink), 6 (cyan), 7 (magenta), 8 (blue), 9 (red), 10 (purple), 11 (ice-blue), 14 (olive), 15 (brown), 20 (yellow); B) 1 (magenta), 2 (blue), 4 (yellow), 5 (green), 13 (cyan), 19 (purple); C) 12 (magenta); D) 3 (red), 16 (yellow), 17 (blue); E) 18 (magenta). E2 state ClusPro clusters: F) 0 (yellow), 3 (red), 4 (orange), 5 (blue), 9 (cyan), 14 (green), 15 (brown); G) 2 (yellow); H) 6 (yellow), 7 (red), 10 (orange), 12 (blue); I) 1 (yellow), 8 (red), 13 (orange); J) 11 (red). The TM1/TM2 site directly borders (and also encompasses) the TM2 helix of the PLN-binding site (i.e., TM2/TM6/TM9) and thus it might function as an accessory or intermediate binding site for it.

**Supplementary Figure 5**

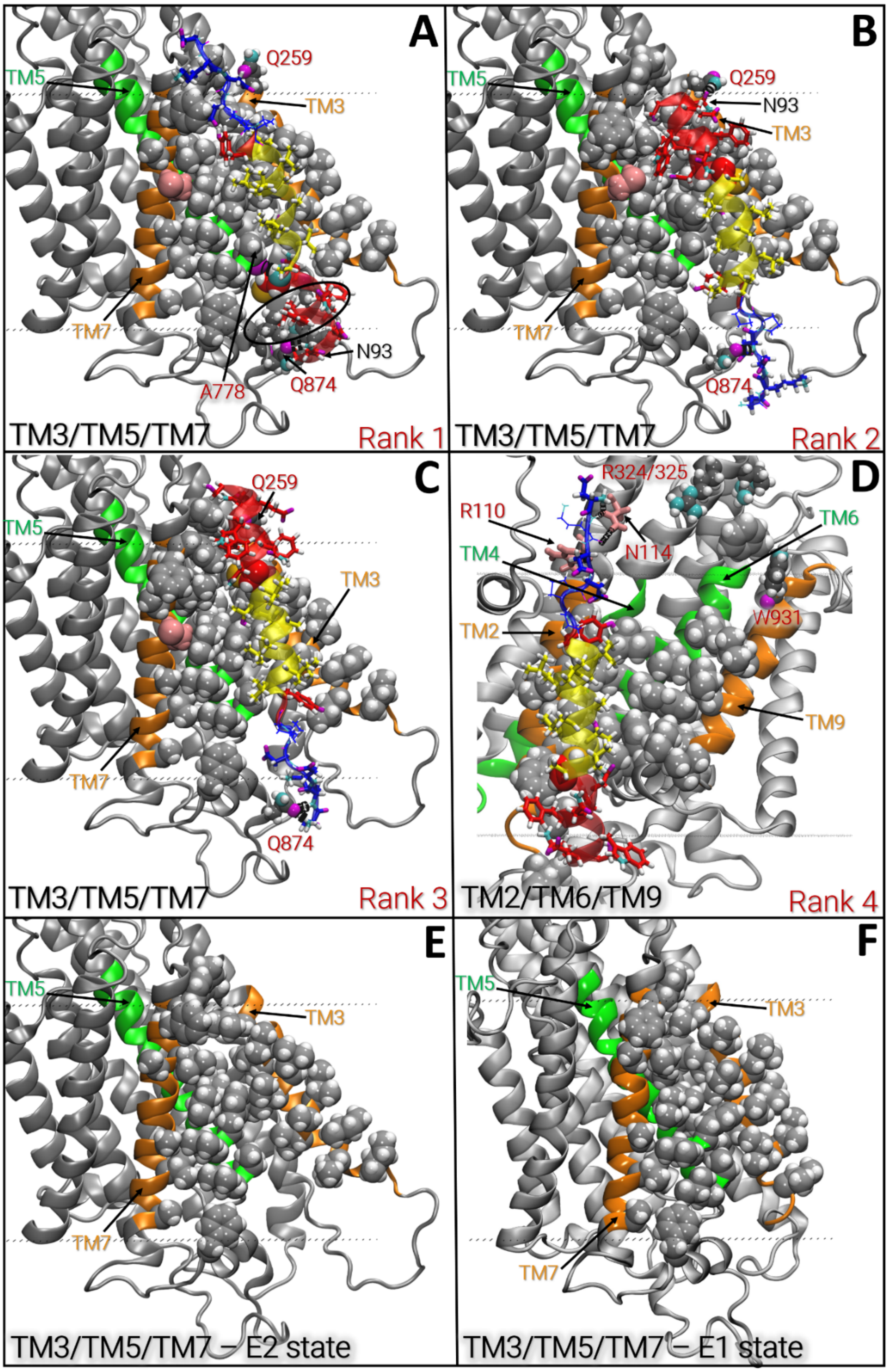

**Supplementary Figure S5 A-F:** Best ranking S100A1ct/SERCA2a models obtained for the E2 state with the MPDock refinement pipeline. Representations and color coding are the same as in Figure 2A (if not mentioned differently, e.g., C841 is shown as pink van der Waals spheres and residues in the tag can be shown as lines). Rank 1 (A), rank 2 (B), and rank 3 (C) all correspond to the TM3/TM5/TM7 docking site. Rank 4 corresponds to the TM2/TM6/TM9 dockings site. Analogously to the E1 state, the large patch of apolar residues shown (here as van der Waals spheres; carbon: gray, hydrogen: white) was selected based on visual inspection of the complex models. Potential polar contacts that can be formed in the complex models according to PyMOL are the following: A) between backbone atoms of C86/S100A1ct and A778/SERCA2a and between the side chains of N93/S100A1ct and Q874/SERCA2a (smaller spheres with dark gray carbon atoms; same representation for most other SERCA2a residues potentially participating in polar contacts) as well as around the area highlighted by a black circle (i.e., between backbone/side chain atoms of W91/S100A1ct and backbone atoms of G291/SERCA2a and W288/SERCA2a and between the side chain of S94/S100A1ct and the backbone of L873/SERCA2a; these are not indicated via black dashes due to compactness of the respective region), B) between the side chains of N93/S100A1ct and Q259/SERCA2a as well as Q874/SERCA2a and residues in the hydrophilic tag of S100A1ct, C) between the side chains of N93/S100A1ct and Q259/SERCA2a as well as Q874/SERCA2a and residues in the hydrophilic tag of S100A1ct. D) The best-ranking TM2/TM6/TM9 docking pose of the E2 state forms potential polar contacts between the SERCA2a residues R110 and N114 (depicted as pink sticks) and S100A1ct residues in the hydrophilic tag, but, contrary to the corresponding docking pose obtained for the E1 state, no polar contacts can be formed between R324/SERCA2a and R325/SERCA2a or W931/SERCA2a and S100A1ct. Interestingly, the large apolar patch of residues at this site seems to be less compact compared to the E1 state. Furthermore, the refined pose is shifted quite substantially within the TM2/TM6/TM9 site compared to the initial ClusPro pose (ClusPro cluster 3 shown in red in Supplementary Figure S4F), while the corresponding refined pose docking to the TM2/TM6/TM9 site in the E1 state (i.e., rank 2) is not substantially moved away from its initial position (refined pose shown in Figure 2A, upper row vs pink ClusPro pose shown in Supplementary Figure S4A). This could indicate that it might be “unhappy” at this site in the E2 state. E) and F): comparison between the compactness of the patch of corresponding apolar residues found at the TM3/TM5/TM7 site of the E2 (E) and E1 (F) states (for both states the SERCA2a side-chain conformations obtained for the best-ranking docking at this site are shown).

**Supplementary Figure 6**

**S6A**

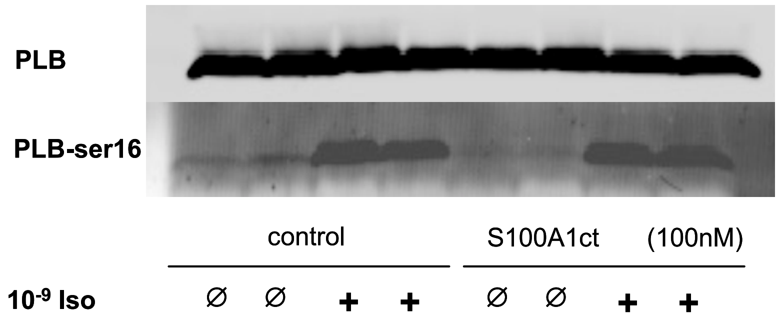

**S6B**

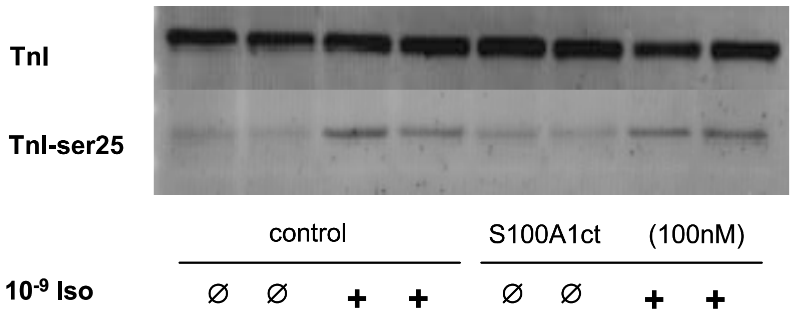

**Supplementary Figure S6 A-B:** Similar increase in phospholamban serine 16 (PLB-ser16; A) and troponin serine 25 (TnI-ser25; B) phosphorylation by the ß-adrenergic receptor mediated agonist isoproterenol ($10^{-9}$M) in cultured AVCMs under control conditions (vehicle) and in the presence of 100 nM S100A1ct peptide, as shown by representative immunoblottings from cellular homogenates for total PLB and TnI and PLB-ser16 and TnI-ser25. Vehicle: aqueous carrier used for S100A1ct peptide dilution.

**Supplementary Figure 7**

**S7**

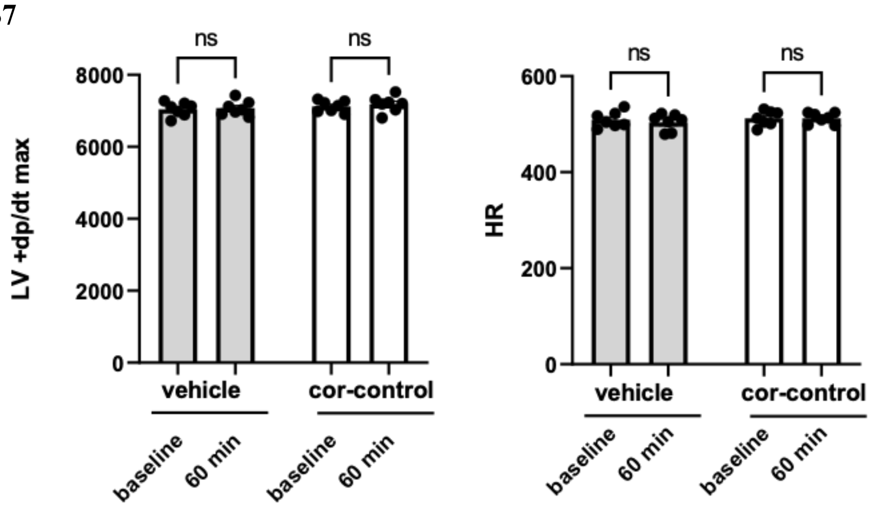

**Supplementary Figure S7:** Neither vehicle nor cor-scrambled control peptide caused an alteration in cardiac contractile performance (left) or heart rate (rate) in healthy mice over the 60 min cor-S100A1ct treatment period. Data are given as mean±SEM.

**Supplementary Table S3**

| Parameter | Reference | control | cor-S100A1ct | P |
|---|---|---|---|---|
| Sodium, mM | (143-148) | 148 | 144 | ns |
| Potassium, mM | (3.5-4.1) | 3.8 | 3.9 | ns |
| Calcium, mM | (2.2-2.4) | 2.3 | 2.3 | ns |
| Albumin, g/l | (25-28) | 26 | 26 | ns |
| Urea, mM | (9.8-12.0) | 10.9 | 11.2 | ns |
| Cholesterol, mM | (2.3-2.9) | 2.6 | 2.5 | ns |
| Triglyceride, mM | (1.4-2.3) | 1.7 | 1.8 | ns |
| Glucose, mg/dl | (140-160) | 144 | 152 | ns |
| GPT, U/l | (24-40) | 32 | 29 | ns |
| GOT, U/l | (40-60) | 51 | 48 | ns |
| AP, U/l | (80-100) | 92 | 91 | ns |
| α-amylase, U/l | (512-618) | 556 | 599 | ns |
| CK, U/l | (550-700) | 720* | 640 | ns |
| hsTnT | (<14pg/ml) | 97* | 23* | <0.05 |
| CRP, mg/dl | | 1.98 | 2.68 | ns |
| White blood cells | ($10^3$/µl) | 3.4 | 3.1 | ns |
| Red blood cells | ($10^6$/µl) | 10.7 | 11.0 | ns |
| Platelet count | ($10^3$/µl) | 1350 | 1402 | ns |
| Neutrophil count | ($10^3$/µl) | 1.2 | 1.4 | ns |
| Lymphocyte count | ($10^3$/µl) | 1.8 | 1.9 | ns |

**Supplementary Table S3:** Clinical chemistry and basic blood count in control (cor-scrambled peptide) and cor-S100A1ct treated mice from 12-day post-treatment blood samples yielded normal organ biomarker and blood count values but significantly greater hsTnT values in the control group (control n=9 vs. cor-S100A1ct n=14 animals P<0.05).

**Supplementary Figure 8**

**S8**

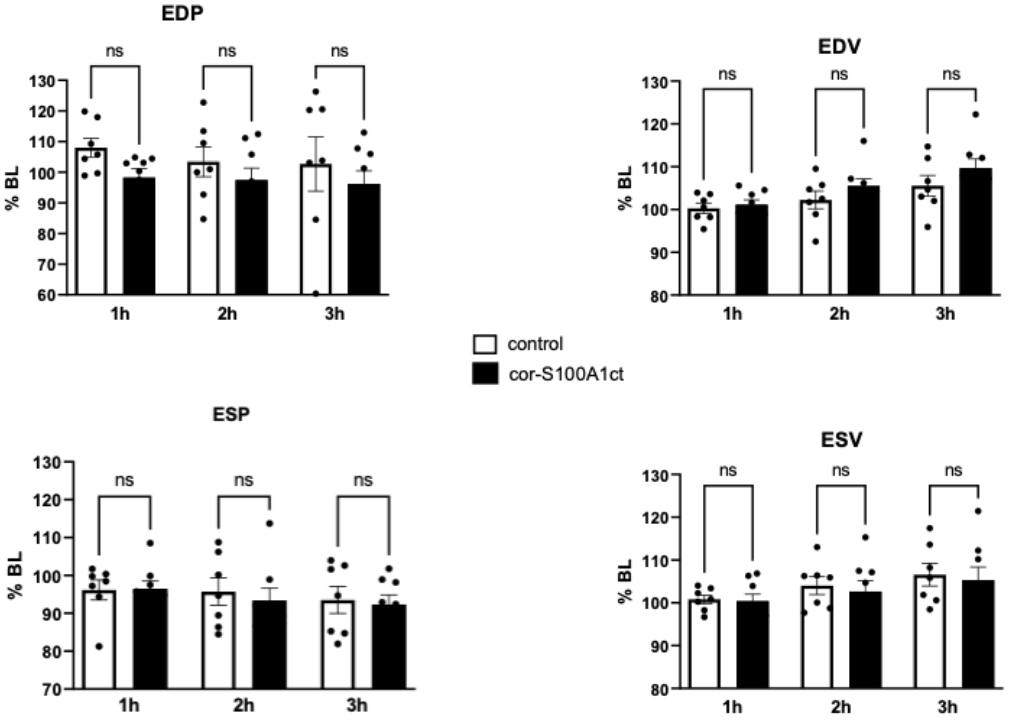

**Supplementary Figure S8:** Neither left ventricular end-diastolic pressure (EDP) nor end-diastolic volume (EDV), end-systolic pressure (ESP) and end-systolic volume (ESV) were different in control (vehicle) and cor-S100A1ct treated pigs during the 3-hour observation period. Data are given as mean±SEM as relative (%) change from individual hemodynamic baseline data. Control n=7; cor-S100A1ct n=8.

**Supplementary Table S4**

| Parameter | Unit | Reference | vehicle | | | S100A1ct | | |
|---|---|---|---|---|---|---|---|---|
| | | | mean | +/- sem | n | mean | +/- sem | n |
| leucocytes | /nl | 10.0-22.0 | 18.86 | 1.38 | 7 | 22.34 | 2.46 | 8 |
| erythrocytes | /pl | 5.8-8.1 | 4.66 | 0.18 | 7 | 5.23 | 0.13 | 8 |
| hemoglobin | g/dl | 10.8-14.8 | 7.56 | 0.30 | 7 | 8.24 | 0.18 | 8 |
| hematocrit | l/l | 0.33-0.45 | 0.23 | 0.01 | 7 | 0.26 | 0.01 | 8 |
| MCV | fl | | 49.57 | 0.57 | 7 | 49.50 | 1.05 | 8 |
| MCH | pg/cell | | 16.43 | 0.20 | 7 | 15.88 | 0.30 | 8 |
| MCHC | g/dl | | 33.29 | 0.52 | 7 | 32.00 | 0.38 | 8 |
| RDW | % | | 16.71 | 0.22 | 7 | 16.38 | 0.34 | 8 |
| platelets | /nl | 175-580 | 266.3 | 19.18 | 7 | 367.50 | 28.67 | 8 |
| sodium | mmol/l | 140-160 | 136.0 | 2.23 | 7 | 135.60 | 1.83 | 8 |
| potassium | mmol/l | 3.5-4.5 | 4.67 | 0.19 | 7 | 4.63 | 0.16 | 8 |
| creatinine | mg/dl | 0.45-1.47 | 0.81 | 0.03 | 7 | 0.95 | 0.06 | 8 |
| urea | mg/dl | 59.4-149.4 | 15.29 | 1.02 | 7 | 16.75 | 1.10 | 8 |
| glucose | mg/dl | 70.2-115.2 | 78 | 5.97 | 7 | 94.50 | 9.21 | 8 |
| hs-troponin T | pg/ml | | 16.43 | 3.23 | 7 | 23.13 | 5.55 | 8 |
| LDH | U/l | <545 | 391.1 | 24.09 | 7 | 495.60 | 49.21 | 8 |
| GOT/AST | U/l | <80 | 34.00 | 10.20 | 7 | 55.00 | 19.36 | 8 |
| GPT/ALT | U/l | <126 | 45.14 | 1.32 | 7 | 44.75 | 2.60 | 8 |
| GGT | U/l | <79 | 26.57 | 1.96 | 7 | 33.38 | 3.25 | 8 |
| lipase | U/l | - | 12.43 | 0.65 | 7 | 11.75 | 0.25 | 8 |
| albumin | g/l | 18-31 | 22.94 | 1.14 | 7 | 24.59 | 1.03 | 8 |
| Quick | % | | 69.63 | 3.08 | 7 | 82.59 | 7.78 | 8 |
| aPTT | s | | 120 | 0.00 | 7 | 65.81 | 20.49 | 8 |

**Supplementary Table S4:** Clinical chemistry and basic blood count in control (vehicle) and cor-S100A1ct treated pigs yielded no abnormal organ biomarker and blood count values after the 3-hour intervention. aPTT showed the expected prolongation due to heparin treatment.
